# Humanized glioblastoma patient-derived orthotopic xenografts recreate a locally immunosuppressed human immune ecosystem amenable to immunotherapeutic modulation

**DOI:** 10.1101/2025.11.20.689484

**Authors:** Pilar M. Moreno-Sanchez, Anaïs Oudin, Batuhan Kisakol, Heiko Dussmann, Eliane Klein, Virginie Baus, Camille Rolin, Carole Seguin-Devaux, Mahsa Rezaeipour, Aurélie Poli, Alessandro Michelucci, Jérôme Paggetti, Etienne Moussay, Jochen H.M. Prehn, Simone P. Niclou, Anna Golebiewska

**Author notes:** Equal contribution (Pilar M. Moreno-Sanchez, Anaïs Oudin). Correspondence should be addressed to A.G.

## Abstract

Immune-based strategies have so far failed to demonstrate clinical benefit in glioblastoma (GBM), largely due to the profound immunosuppressive tumor microenvironment (TME). To achieve more predictive preclinical insights, advanced *in vivo* models that faithfully recapitulate the human brain immune landscape are urgently needed. Here, we established GBM patient-derived orthotopic xenografts (PDOXs) across diverse mouse strains, including humanized models. Humanization was achieved through transplantation of CD34^+^ hematopoietic stem cells (HU-CD34^+^) or peripheral blood mononuclear cells (HU-PBMC). Both models successfully reconstituted human T-cells systemically, with stronger engraftment in HU-CD34^+^ mice. We observed selective infiltration and spatial organization to intracranial GBM tumors, including exhausted, memory-like, and regulatory CD4^+^ T-cell phenotypes, TIM-3^+^ immunosuppressive-like myeloid cells and intratumoral B cells. Mouse microglia-derived tumor-associated macrophages (TAMs) remained the dominant immunosuppressive immune population. Anti-PD-1 therapy, but not anti-GITR, modestly modulated the infiltration dynamics, demonstrating the susceptibility of the reconstructed adaptive immunity to immunotherapeutic intervention. These findings position humanized GBM PDOXs as a relevant preclinical platform to interrogate tumor-immune interactions and evaluate immunotherapeutic strategies in a human context.

**Key points:**

- GBM PDOXs developed in HU-CD34^+^ and HU-PBMC mice faithfully reconstitute systemic and local human adaptive immunity.
- Human immune components undergo selective infiltration, spatial organization and transition towards exhausted CD4^+^ T-cells and immunosuppressive CD11c^+^ myeloid cells.
- Anti-PD-1, but not anti-GITR, locally promote human immune infiltration into intracranial GBM tumors, while sparing systemic compartments.
- Humanized GBM PDOXs provide a powerful preclinical platform to test novel immunotherapeutic strategies.

**Study importance:** Immune checkpoint blockade has shown limited efficacy in GBM, reflecting the highly immunosuppressive and lymphocyte-poor nature of the TME. Conventional syngeneic and GEMM models fail to recapitulate these features, contributing to the translational disconnect between preclinical success and clinical failure. Humanized mice provide a solution to interrogate human-specific immunity *in vivo*, but their use in GBM has remained limited. Here, we provide the first comparison of GBM PDOX modeling in two complementary modes of humanization based on CD34^+^ HSCs and PBMCs. We systematically profile systemic and intratumoral compartments, showing that these models faithfully reconstitute human adaptive immunity and capture the interplay with the murine brain TME. Furthermore, we demonstrate clinically-relevant responses upon treatment with checkpoint antibodies targeting PD-1 and GITR, showing modulation of human immune subsets without altering murine TAM immunosuppression, underscoring the translational value of the system. This study establishes humanized GBM PDOXs as a versatile platform for dissecting tumor-immune interactions in the brain and for preclinical evaluation and development of novel immunotherapies.

**Graphical abstract:** 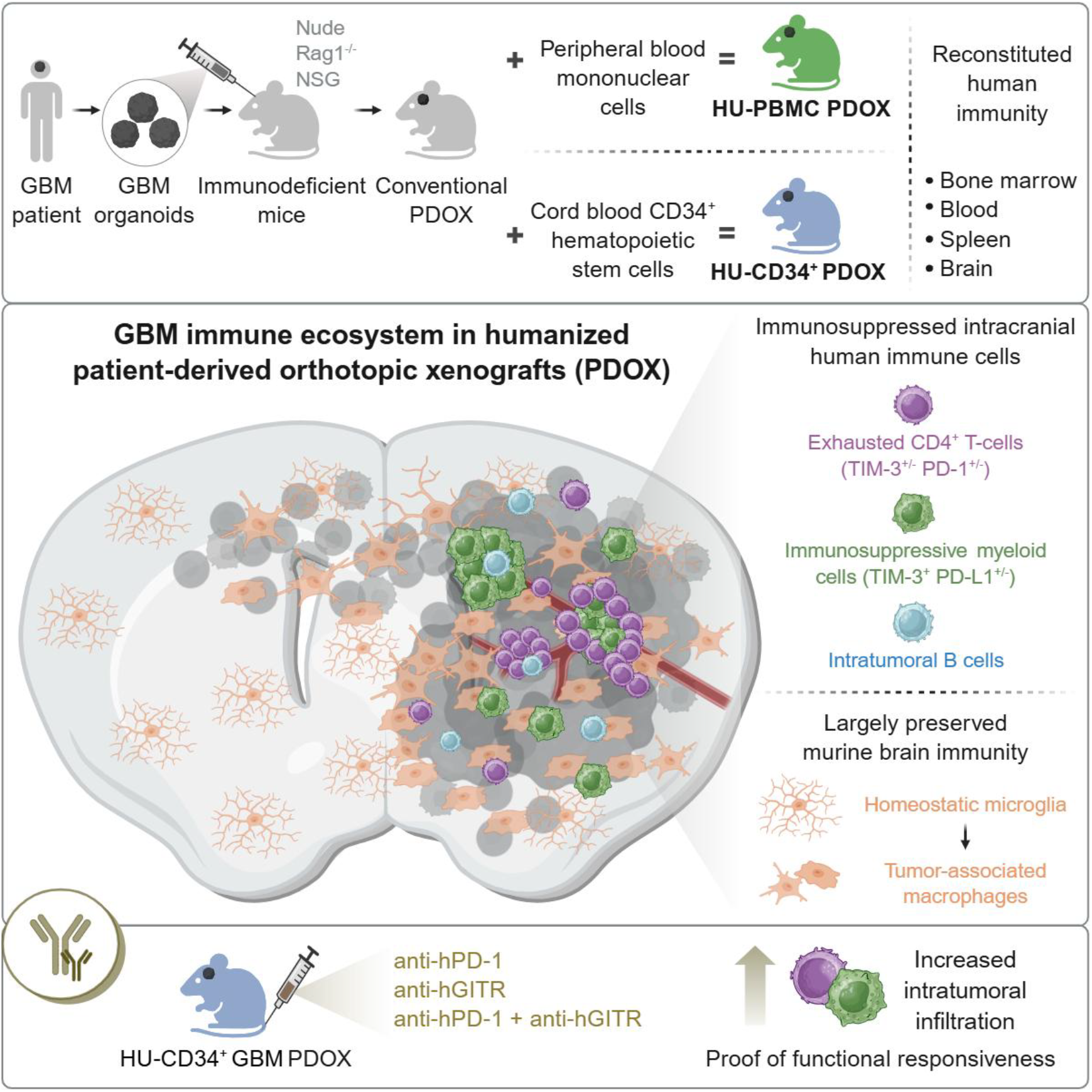

## Introduction

Glioblastoma (GBM) remains the most aggressive primary brain tumor, with rapid progression and poor prognosis despite multimodal standard-of-care treatment^1^. Immunotherapy has transformed outcomes in other solid tumors, yet its impact in GBM has been minimal^2^. This lack of efficacy reflects the profoundly immunosuppressive and “cold” GBM tumor microenvironment (TME), characterized by abundant tumor-associated macrophages (TAMs) with tolerogenic functions and sparse T-cell infiltration, enriched for regulatory or exhausted CD4^+^ T-cells with effector memory phenotypes^2-4^. Preclinical studies of immune checkpoint blockade (ICB) in GBM have traditionally relied on syngeneic or genetically engineered mouse models (GEMMs). While these immunocompetent systems allow mechanistic interrogation, they fail to capture the molecular heterogeneity of patient tumors and the GBM-specific immunosuppressive milieu, often yielding false-positive responses^5-7^. Consistently, multiple clinical trials, including those testing anti-PD-1 antibodies, have not demonstrated survival benefit in patients with GBM^8-10^, underscoring the translational gap between conventional models and the clinic. A growing consensus suggests that therapeutic efficacy in GBM will require rational combinations, first ‘priming’ the TME to overcome myeloid-driven suppression, and subsequently ‘rewiring’ adaptive immunity to sustain effector function^11^. Humanized mice, generated by engrafting human hematopoietic progenitors (HU-CD34^+^) or peripheral blood mononuclear cells (HUPBMCs), offer an avenue to bridge the preclinical gap. These systems reconstitute functional human immune compartments and have proven invaluable for modeling human-specific immunity in contexts such as autoimmunity^12,13^ and infectious diseases, particularly HIV^14-17^. Their application in cancer remains relatively novel, but they hold promise for interrogating the interplay between human tumor and immune compartments and for testing immunotherapies in a more patient-relevant setting.

To exploit this potential in GBM, we developed patient-derived orthotopic xenografts (PDOXs) in humanized mice. We previously showed that PDOXs derived from GBM organoids preserve patient-specific histopathology and molecular features while engaging in reciprocal crosstalk with the murine brain TME, including vasculature and microglia-derived TAMs^18-20^. We have shown that these murine-derived components are functionally responsive to therapy, including surgery^21^, temozolomid^20^, and anti-angiogenic agents^22^. Here, we extend this strategy by combining GBM PDOXs with two modes of humanization based on HU-CD34^+^ and HU-PBMCs mice, and directly comparing their capacity to reconstitute systemic and intratumoral immunity. We show that GBM tumors reliably develop in humanized mice, and we provide a comprehensive phenotypic and functional profiling of human and murine immune cells across the blood, spleen, bone marrow, and brain in tumor-bearing and healthy counterparts. These models successfully reconstitute a human adaptive immune system both systemically and intratumorally, offering a closer approximation to the immune landscape observed in GBM patients. We further tested systemic administration of anti-PD-1 and anti-GITR antibodies, showing modest shifts in human immune composition and T-cell functional states, without affecting murine TAM immunosuppressive features. Together, these results establish humanized GBM PDOXs as a translationally relevant platform to test immunotherapies, bridging a critical gap between conventional models and clinical trials.

## Results

### Baseline brain immune landscape is preserved among immunocompromised mouse strains

Systemic immune components vary across different mouse strains, influencing the likelihood of human tumor cell engraftment (**Fig. 1A**). We have derived a cohort of >45 glioma PDOX models by implanting patient-derived organoids in NSG mice^18^. Established models can also efficiently engraft in Nude^20^, and Rag1KO mice. While the peripheral immune system of these strains has been extensively characterized^23-25^, differences in brain immunity remain underexplored. We observed no differences in anatomical or structural integrity in healthy brains of three immunodeficient strains (Nude, Rag1KO, NSG) compared to C57BL/6J immunocompetent mice (**Fig. S1A**). Iba1^+^ myeloid cells displayed a similar distribution across white and grey matter tracts in different brain regions (**Fig. S1B**), consistently exhibiting ramified morphology of homeostatic microglia (Mg)^26^. Dominance of Mg over monocyte-derived macrophages (Mo) and border-associated macrophages (BAMs) was consistent across strains (**Fig. 1B, S1C)**, with no significant differences in CD45 and CD11c expression **(Fig. S1D**), suggesting no major differences in their activation status. Lymphocytes were rare within the brain, with minor variations reflecting different levels of immunocompetency across strains **(Fig. 1B**). Overall, these findings indicate that immunodeficiency does not fundamentally impact the baseline brain immune environment.

**Figure 1.**
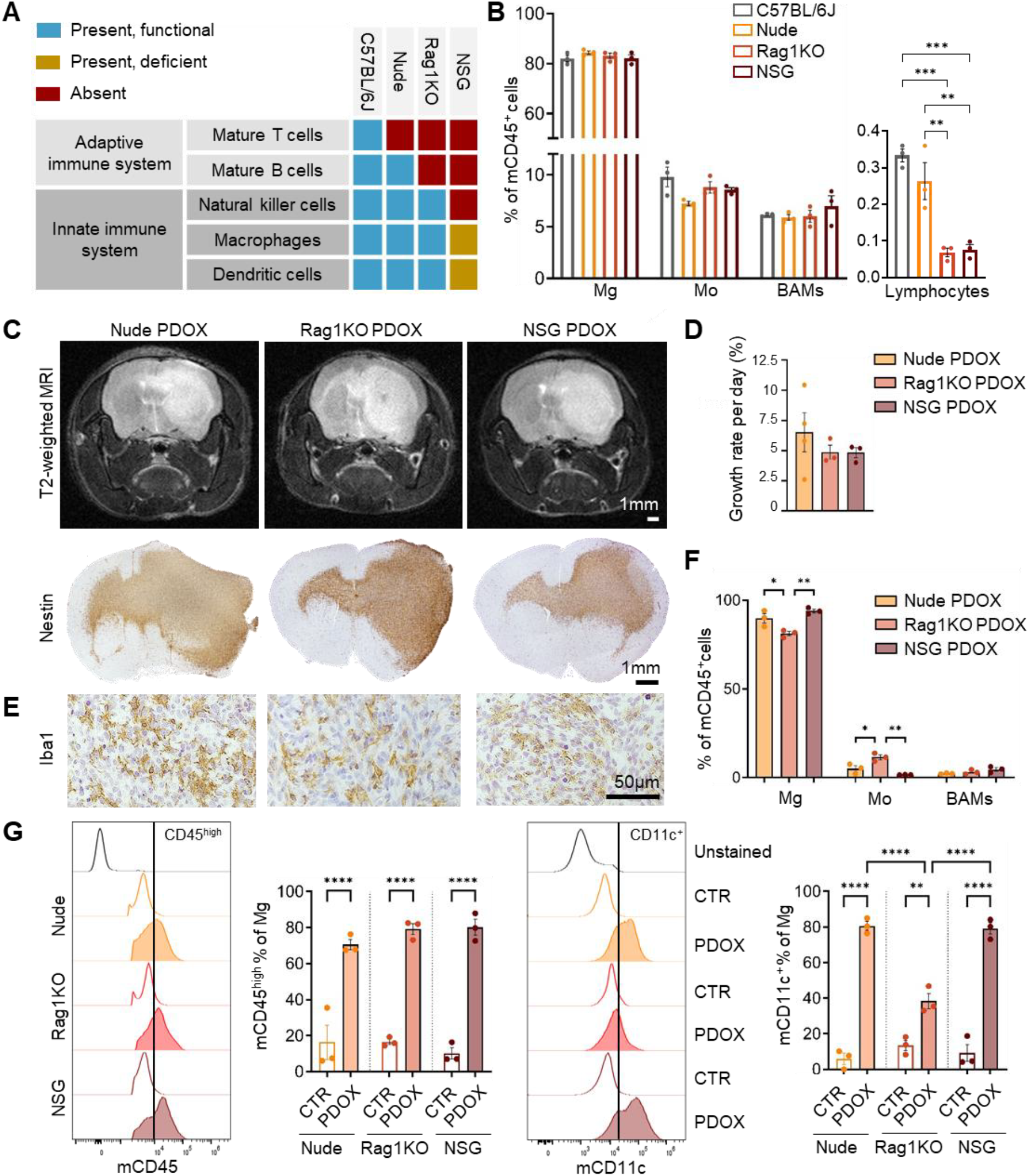
Immune profiling of brains in diverse mouse strains at baseline and GBM PDOXs. **A.** Schematic summarizing the systemic immune components in diverse mouse strains. **B**. Flow cytometry quantification of microglia (Mg), monocyte-derived macrophages (Mo), border-associated macrophages (BAMs) and lymphocytes in brains of diverse mouse strains; one-way ANOVA with Tukey’s HSD correction per each cell type (n=3 mice/strain, mean ± SEM, **p<0.01, ***p<0.001). See gating strategy in **Fig. S1C. C**. Representative MRI and immunohistochemistry images of coronal sections in entire brains of PDOX T188 developed in Nude, Rag1KO and NSG mice. Nestin depicts human GBM cells; scale bar: 1 mm. **D**. Average growth rate of PDOX T188 per day in three mouse strains between the tumor detection and endpoint; one-way ANOVA with Tukey’s HSD correction (n = 4 mice in Nude, n = 3 mice in Rag1KO and NSG, mean ± SEM, ns = not significant). **E**. Iba1 staining showing ameboid Mg within the tumor core across all strains. Images show magnified views of cellular tumor areas; scale bar: 50 μm. **F**. Flow cytometry quantification of the Mg, Mo and BAMs within CD45^+^ immune cells in PDOX T188 developed in different stains (n=3 mice/strain, mean ± SEM, one-way ANOVA with Tukey’s HSD correction, *p<0.05, **p<0.01). See **Fig. S1C** for gating strategy. **G**. Representative flow cytometry histograms and quantification showing increased CD45 and CD11c expression in Mg within tumors of PDOX T188-implanted mouse brains compared to healthy control brains (CTR) across strains; one-way ANOVA with Tukey’s HSD correction (n=3 mice/condition, mean ± SEM, **p<0.01, ****p<0.0001).

### Functional crosstalk between human GBM cells and mouse Mg is maintained in PDOXs across diverse immunocompromised mouse strains

We next assessed whether human GBM cells engage in similar functional crosstalk with mouse Mg across PDOXs from different strains. We previously demonstrated that TAMs in PDOXs derived in Nude mice arise largely from mouse-derived Mg carrying dendritic cell-like (DC) signatures, while Mo and BAMs show lower contribution to TAMs^20^. To investigate whether Mg can functionally interact in other immunodeficient mouse stains, we established GBM PDOXs in Nude, Rag1KO, and NSG mice. We selected four representative PDOX models with different histopathological features and molecular background^18,27^ (**Table S1**). MRI and histopathological analyses confirmed comparable tumor volumes and growth rates across the strains (**Fig. 1C-D, S2A)**. PDOXs exhibited comparable behavior across strains, retaining their invasive (P8), intermediate (T188, P3), or angiogenic (P13) histopathological growth patterns. Notably, angiogenic PDOXs developed pseudopalisade-like structures characteristic of GBM in all strains (**Fig. S2A**). Iba1 staining revealed ameboid-shaped TAMs within tumor cores, indicative of GBM-specific education, with comparable density and spatial distribution in all mouse strains (**Fig. 1E, S2A**). Mg remained the predominant source of TAMs, where only a minor increase in Mo was observed in PDOXs in Rag1KO mice (**Fig. 1F**). Mg-TAMs displayed elevated CD45 expression relative to healthy brain controls across all strains (**Fig. 1G**), further supporting GBM-specific activation. Interestingly, Mg, Mo and BAMs in tumors derived in Rag1KO mice exhibited lower CD11c expression to that observed in Nude and NSG PDOXs (**Fig. 1G, S2B-C**), suggesting a reduced activation state of myeloid cells or diminished interaction with human GBM cells in this strain. This is consistent with a recent study reporting less reactive Mg in Rag1KO mice compared to C57BL/6J in a brain ischemia model^28^. Collectively, these results indicate that GBM PDOX tumor growth and murine TAM features in PDOXs are preserved across diverse immunodeficient strains. Since transition of Mg towards CD11c^+^ TAMs in Rag1KO mice is less pronounced, this strain may be less suitable for studies requiring robust TAM activation.

### HU-CD34^+^ and HU-PBMC humanized mouse models support successful intracranial growth of GBM PDOX tumors

We further advanced our *in vivo* model by introducing a human adaptive immune system in a NSG mouse background. We compared two humanization approaches: HU-CD34^+^ mice, engrafted with CD34^+^ cord blood HSCs; and HU-PBMC mice, engrafted with mature PBMCs (**Fig. 2A**). The two humanization protocols required different strategies for induction of tumor growth^29^. HU-CD34^+^ model requires depletion of mouse immune cells through irradiation, with humanization confirmed as >25% human CD45^+^ cells in the blood at 12-15 weeks after HSC engraftment, followed by intracranial implantation of GBM organoids. In HU-PBMC mice, GBM organoids were implanted before intravenous PBMC transfer, as these mice rapidly restore mature T-cells in the circulation but develop GvHD within 3–4 weeks after humanization. We selected two PDOX models, T188 and T158, based on the HLA profiles and similar characteristics (**Table S1**). Gamma-irradiated HU-CD34^+^ was exclusively applied for PDOX T188, whereas busulfan-treated HU-CD34^+^ and HU-PBMC were applied for both T188 and T158. In the gamma-irradiated HU-CD34^+^ model, donor immune cells were matched to PDOX T188 for the HLA-A loci (**Table S2**). For the busulfan-irradiated CD34^+^ model, no information on donor HLA loci was available. In HU-PBMC model, several HLA loci aligned with T188 and T158.

**Figure 2.**
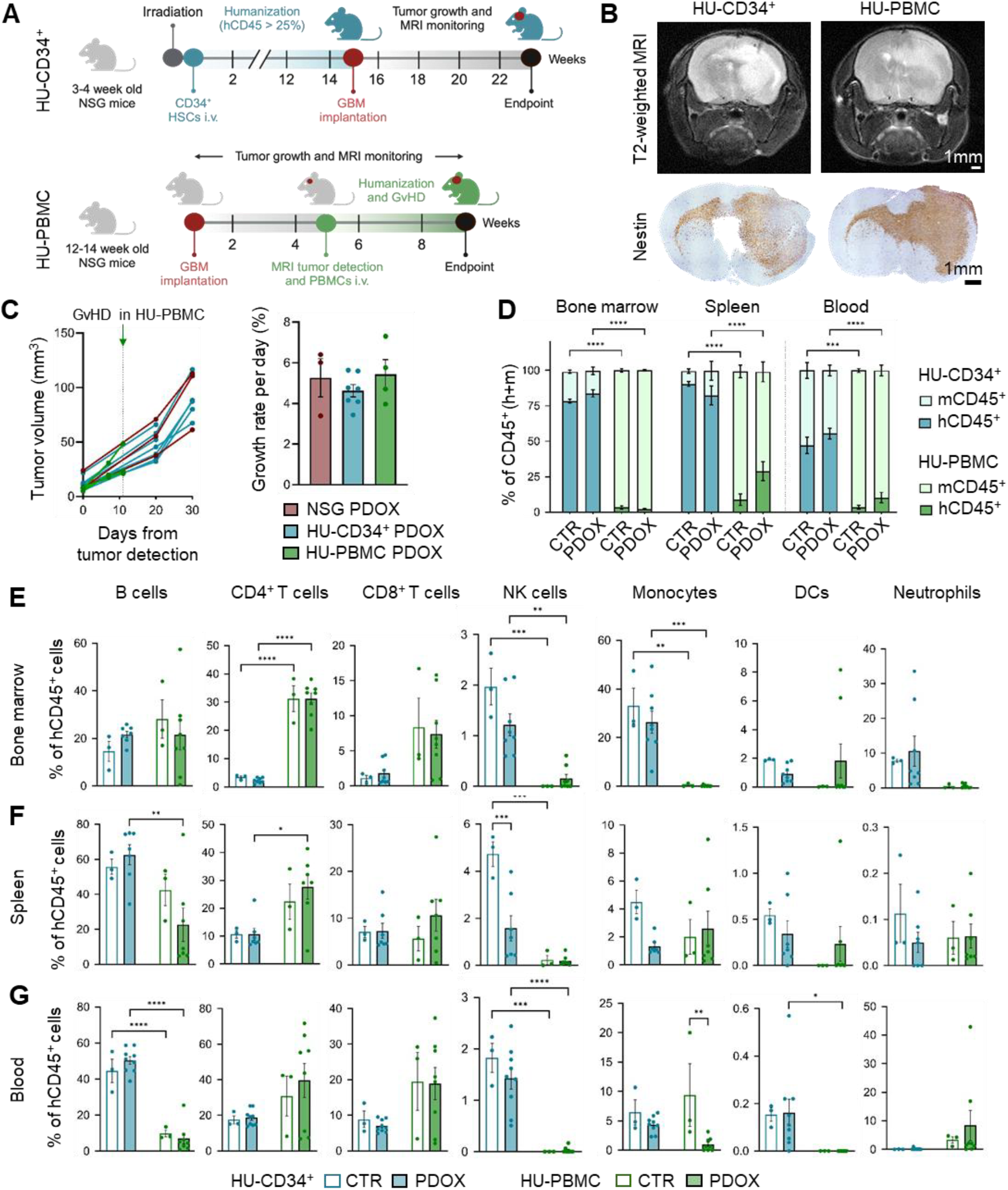
HU-CD34^+^ and HU-PBMC PDOXs show efficient tumor growth and systemic reconstitution of human immune cells. **A.** Schematic illustrating key steps in generation of GBM PDOXs in HU-CD34^+^ and HU-PBMC mice. **B**. Representative MRI and immunohistochemistry images of coronal sections of the entire brains in PDOX T188 developed in humanized mice. Nestin depicts human GBM cells; scale bar: 1 mm. **C**. MRI-based tumor volume (mm^3^) and growth rate quantification in PDOX T188 developed in NSG, HU-CD34^+^ and HU-PBMC mice. Each connected dot line represents an individual mouse; one-way ANOVA with Tukey’s HSD correction (n=3 for NSG and n=4-7 for PDOX mice/condition, mean ± SEM, ns = not significant). **D**. Ratio between human (hCD45^+^) and mouse (mCD45^+^) immune cells in bone marrow, spleen, and blood in HU-CD34^+^ (irradiated and busulfan combined) and HU-PBMC control mice (CTR) and PDOX T188; one-way ANOVA with Tukey’s HSD correction (n=3-5 for CTR and n=7-9 for PDOX mice/condition, mean ± SEM, ***p<0.001, ****p<0.0001). See gating strategy in **Fig. S3C. E-G**. Flow cytometry quantification of human lymphoid (B, T, NK cells) and myeloid populations (monocytes, DCs, neutrophils) in bone marrow (E), spleen (F), and blood (G) in HU-CD34^+^ and HU-PBMC CTR and PDOX T188 mice; one-way ANOVA with Tukey’s HSD correction (n=3 for CTR and n=7-9 for PDOX mice/condition, mean ± SEM, *p<0.05, **p<0.01, ***p<0.001, ****p<0.0001).

Weekly MRI revealed consistent engraftment and tumor growth of GBM PDOXs T188 (**Fig. 2B**) and T158 (**Fig. S3A**) in HU-CD34^+^ and HU-PBMC mice. Tumor growth and histopathological features remained similar, comparable to NSG mice (**Fig. 2C, S3B**), suggesting that human immune cells, irrespective of HLA compatibility, do not hinder tumor growth in humanized PDOXs. HU-CD34^+^ mice were euthanized at the humane endpoint due to neurological symptoms associated with the intracranial tumor. HU-PBMC mice were sacrificed at 3-4 weeks post-PBMC injection to prevent GvHD^30^. Only one HU-PBMC mouse exhibited GvHD-associated symptoms, including fur loss, ruffled coat, hunching, and significant weight loss. The remaining HU-PBMC mice displayed tumor-associated neurological symptoms. These findings confirm that humanized mice support successful derivation of GBM PDOXs, providing an advanced *in vivo* platform for studying adaptive immunity in GBM.

### HU-CD34^+^ mice develop higher systemic humanization levels compared to HU-PBMC mice, amenable to tumor-induced modulation

We further investigated the extent of systemic humanization in bone marrow, spleen and blood at the endpoint of tumor development (**Fig. 2D, S3C-D**). Overall, HU-CD34^+^ mice demonstrated significantly higher proportions of human CD45^+^ immune cells in all anatomical sites compared to HU-PBMC mice in both healthy control mice (CTR) and tumor-bearing PDOXs (**Fig. 2D**). These differences were expected since PBMCs do not require bone marrow engraftment. In contrast, CD34^+^ HSCs rely on bone marrow engraftment for sustained *in situ* maturation and immune reconstitution^31^. PBMCs have a limited lifespan in circulation, leading to less sustained immune cell reconstitution compared to CD34^+^ models. Proportions of human CD45^+^ cells were higher in bone marrow and spleen of HU-CD34^+^ mice compared to blood. Unlike NSG mice, HU-CD34^+^ mice exhibited restored splenic white pulp and integration of human CD45^+^ cells (**Fig. S3E, S3F-G**), indicating partial reconstitution of adaptive immunity^32-34^. Interestingly, both HU-CD34^+^ and HU-PBMC PDOXs exhibited enlarged spleens and increased proportions of human CD45^+^ cells (**Fig. S3H, S3I**) in comparison to their healthy counterparts (**Fig. 2D, S3D**), indicating tumor-induced immune activation (**Fig. S3H**). These findings also extend to HUPBMC mice (**Fig. S3I**) with increased proportions of human CD45^+^ cells compared to their respective controls (**Fig. 2D, S3D**). This observation further demonstrates that intracranial GBM tumors can significantly modulate the systemic immune system, highlighting brain-systemic communication, consistent with observations in human patients. Overall, our data shows higher levels of humanization in HU-CD34^+^ versus HU-PBMC mice, retained in both CTR and tumor-bearing humanized mice.

### HU-CD34^+^ and HU-PBMC mice present model-specific systemic humanization composed of lymphocytic and monocytic components

We further applied multicolor flow cytometry (**Fig. S3C**) to evaluate composition of the human lymphoid and myeloid cells across bone marrow, spleen and blood. Both models successfully reconstituted human immune cells in the circulation. Human lymphoid cells dominated over myeloid cells in the spleen and blood in both models, and only HU-CD34+ bone marrow showed a more balanced distribution between lymphocytes and monocytes in the bone marrow (**Fig. 2E-G, S4A-C**). The lymphoid compartment of HU-CD34^+^ mice was primarily composed of B cells, CD4^+^ and CD8^+^ T-cells, and rare NK cells. HU-PBMC mice showed lower proportions of B cells in spleen and blood compared to HU-CD34^+^ mice (**Fig. 2E-G, S4A-C**), with a lymphocytic compartment characterized mainly by CD4^+^ and CD8^+^ T-cells, and low abundance of NK cells. The reconstitution of human myeloid cells was mainly limited to classical monocytes (CD14^+^CD16^-^) and neutrophils (CD66^+^), which were detected at the highest levels in the bone marrow of HU-CD34^+^ mice (**Fig. 2E, S4A**), but also in spleen and blood (**Fig. 3F-G, S4B-C**). DCs (CD11b^+^CD11c^+^HLA^-^DR^+^) were also very low. CD4^+^CD8^+^ double-positive T-cells were visible in the bone marrow and spleen of HU-PBMC mice, but not HU-CD34^+^ mice (**Fig. S4D**). The input PBMCs contained 5.3% of CD4^+^CD8^+^ T-cells and their expansion is known to play a role in the onset of graft-versus-host disease (GvHD)^35^, a known limitation of the HU-PBMC mice. Overall, both models successfully reconstituted human immune cells in the circulation, with the HU-CD34^+^ model demonstrating superior reconstitution of B cells in blood and spleen, while the HUPBMC humanization was limited to T-cells.

**Figure 3.**
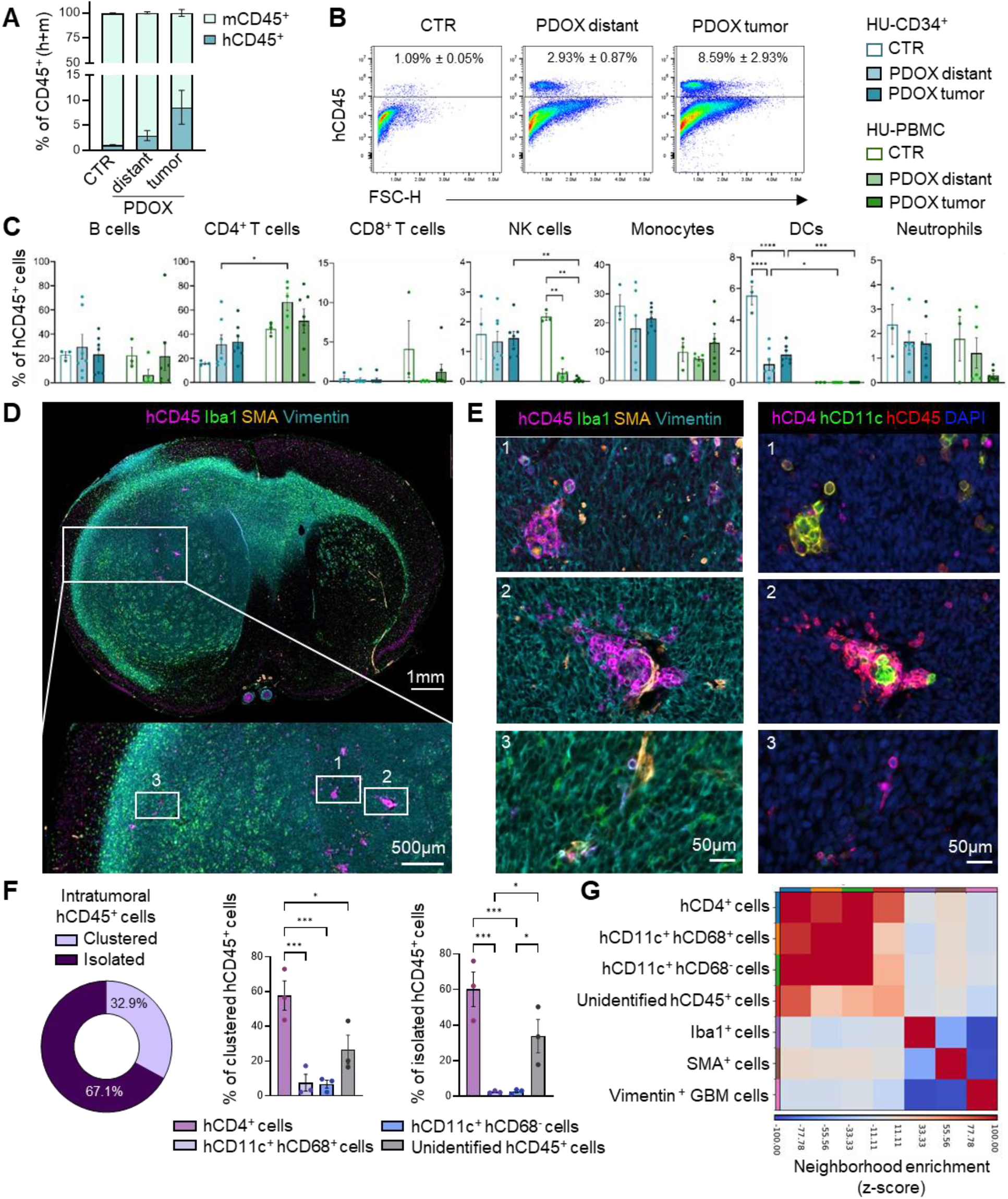
Human immune cells selectively infiltrate GBM tumors in humanized mice and display distinct spatial organization within the TME. **A.** Ratio between human (hCD45^+^) and mouse (mCD45^+^) immune cells in the brains of HU-CD34^+^ control (CTR) mice and PDOX T188 tumor-bearing mice. PDOX brain was divided into cellular tumor (‘tumor’, right tumor-bearing hemisphere) and distant brain (‘distant’, left hemisphere including invasive normal brain areas)**; o**ne-way ANOVA with Tukey’s HSD correction (n=3 for CTR, n=7 for PDOX mice/condition, mean ± SEM, ns = not significant). **B**. Representative flow cytometry dot plots showing hCD45^+^ immune cells in control (CTR), and PDOX T188 “distant” and “tumor” areas of HU-CD34^+^ mice. Mean ± SEM is shown. **C**. Flow cytometry quantification of human lymphoid and myeloid cells in the brains of HU-CD34^+^ and HU-PBMC mice for control (CTR) and PDOX T188 mice (‘tumor’, and ‘distant’ areas); one-way ANOVA with Tukey’s HSD correction (n=3 for CTR and n=5-7 for PDOX mice/condition, mean ± SEM, *p<0.05, **p<0.01, ***p<0.001, ****p<0.0001). See gating strategy in **Fig. S3C. D**. Representative multiplex immunofluorescence of HU-CD34^+^ PDOX T188 brain, highlighting hCD45^+^ immune cells (magenta), Iba1^+^ myeloid cells (green), SMA^+^ blood vessels (orange), and Vimentin^+^ tumor cells (cyan). The large panel shows the overall tumor region (scale bar: 1 mm), with a magnified view of the tumor core (scale bar: 500 µm) illustrating the three regions of interest (ROIs) selected for subsequent high-resolution imaging. ROIs 1 and 2 correspond to intratumoral immune cell clusters, whereas ROI 3 represents isolated human immune cells located toward the tumor periphery. **E**. High-magnification multiplex immunofluorescence of the three ROIs shown in (D). The first panel displays human CD45^+^ immune cells (magenta), Iba1^+^ myeloid cells (green), SMA^+^ blood vessels (orange), and Vimentin^+^ tumor cells (cyan), while the second panel shows staining for hCD4 (pink), hCD11c (green), hCD45 (red), and DAPI (blue). Scale bar: 50 µm. **F**. Quantification of intratumoral hCD45^+^ immune cells depicting the proportion of clustered versus isolated cells (mean of n=3 biological replicates). Bar plots represent the relative distribution of immune subtypes within each group, one-way ANOVA with Tukey’s HSD correction (n=3, mean ± SEM, *p<0.05, ***p<0.001). **G**. Heatmap of neighborhood enrichment showing spatial proximity among the cell subtypes identified by multiplex imaging. The color scale represents neighborhood enrichment z-scores.

We observed remarkable similarity in the human immune cell composition in HU-CD34^+^ PDOX T188 and T158 and their healthy counterparts (**Fig. 2E-G, S4A-C**). HU-CD34^+^ PDOX mice showed tendency toward elevated proportions of B cells, with statistical differences detected in PDOX T158’s spleen and blood (**Fig. S4B-C**). While T-cell proportions remained similar, we observed a tendency toward general decrease in NK cells and classical monocytes, with statistically significant changes in PDOX T158 (**Fig. S4A-C**). HU-PBMC PDOXs also retained similar immune cell composition, though elevated proportions of human CD45^+^ cells were observed in spleen compared to control mice (**Fig. S3D)**. Yet no specific immune cell type was selectively increased (**Fig. S4B**), reinforcing the hypothesis of tumor-specific activation of immune cells in the spleen. In summary, both HU-CD34^+^ and HU-PBMC mice achieved effective systemic humanization and exhibited model-specific human immune cell composition.

We further evaluated T-cell functionality by assessing activation following *ex vivo* stimulation with PMA/Ionomycin. This was particularly important for HU-CD34^+^ mice, since human T-cells develop entirely within a mouse environment. CD3^+^ T-cells isolated from the spleen of HU-CD34^+^ and HU-PBMC mice exhibited a cytokine profile partially resembling PBMCs (**Fig. S4E-F**). While the degranulation capacity of CD8^+^ T-cells was suboptimal (**Fig. S4E**), the production of IFN-γ, TNF-α, and IL-2 (**Fig. S4E-F**) upon stimulation by both CD4^+^ and CD8^+^ subsets indicates a degree of functional competency. These results suggest that T-cells are not anergic, as they retain cytokine-secreting potential, although their functionality may be limited in other aspects.

### Human CD4^+^ T-cells, B cells and classical monocytes preferentially infiltrate intracranial tumors

We further investigated whether humanized PDOXs reconstituted the local human TME components observed in human GBM. We analyzed the right hemisphere, which contains the primary cellular tumor niche (“PDOX tumor”) and the contralateral left hemisphere, only partially infiltrated by GBM cells (“PDOX distant”) (**Fig. 3A-B**). Brains from healthy humanized mice (CTR) served as controls. Flow cytometry revealed presence of human CD45^+^ cells in PDOX brains, at higher levels in the tumor compared to the distant brain, with minimal immune cell infiltration observed in CTR brains (**Fig. 3A-B**). Human CD45^+^ cells infiltrating tumors and, to a lesser extent, control brains, primarily consisted of CD4^+^ T-cells, alongside B cells and classical monocytes in PDOX T188 (**Fig. 3C**) and T158 (**Fig. S3C, S5A**). Rare NK cells, neutrophils, DCs and CD8^+^ T-cells were also detected. CD4^+^ T-cells were proportionally higher in HU-PBMC compared to HU-CD34^+^ in both PDOX models, whereas higher proportions of monocytic cells and DCs were detected in HU-CD34^+^ brains (**Fig. 3C, S5A**), consistent with model-dependent differences in systemic human immune cell reconstitution (**Fig. 2E-G, S4A-C**). Notably, CD4^+^CD8^+^ double-positive T-cells were nearly absent in HU-PBMC and undetectable in HU-CD34^+^ brains (**Fig. S5B**), confirming that GvHD predominantly manifests as a systemic condition, with limited neurological symptoms^36,37^.

### Intratumoral human CD45^+^ cells show diverse spatial organizational patterns, with frequent clusters of CD4^+^ T-cells and CD11c^+^CD68^+/-^ myeloid cells

To examine distribution of the infiltrating immune cells within the tumor, we performed multiplex immunofluorescence on HU-CD34^+^ T188 PDOX brain sections. We detected Vimentin^+^ GBM cells, Iba1^+^ myeloid cells, and mouse-specific SMA^+^ vasculature (**Fig. 3D**). Within human CD45^+^, we distinguished subsets specified by CD4, CD11c and CD68 positivity, while CD8 and CD19 evaluation was not reliable (**Table S3**). Human CD45^+^ cells were detected within Vimentin^+^ tumor regions, forming large clusters primarily in the central tumor core and smaller clusters or isolated cells scattered across the whole tumor, but not in normal brain regions (**Fig. 3D**). While some clusters were located near SMA^+^ vessels, immune cells also diffused throughout the tumor tissue, independent of visible mature vasculature (**Fig. 3D-E**).

Within Vimentin^+^ tumor regions, approximately 30% of human CD45^+^ cells were organized in heterogeneous aggregates composed predominantly of CD4^+^ T-cells along with CD11c^+^CD68^+/-^ myeloid cells and CD45^+^CD4^−^CD68^−^CD11c^−^ unidentified immune cells (**Fig. 3E-F**). While we were unable to phenotypically resolve this population by imaging, their proportion closely mirrored B cells detected by flow cytometry. CD11c^+^CD68^+^ myeloid subset likely corresponds to Mo-derived DC-like TAMs, whereas CD11c^+^CD68^-^ subset aligns with conventional DCs^38,39^. Myeloid populations showed tendency to accumulate in spatially organized niches (**Fig. S5C-D**), although were also found as isolated single cells (**Fig. 3F**). Neighborhood analyses further confirmed that CD4^+^ T-cells and CD11c^+^CD68^+/-^ myeloid cells preferentially localized in proximity to cells of their own lineage (**Fig. 3G**). Moreover, CD11c^+^CD68^+/-^ myeloid subsets showed a neighborhood enrichment with CD4^+^ T-cells, consistent with direct spatial contacts observed in identified heterogeneous clusters (**Fig. 3E**). Interestingly, some immune clusters showed organized structures, where CD4^+^ cells surrounded CD11c^+^ cells (**Fig. 3E**), reminiscent of the myeloid and T-cell dyads described in human gliomas^40^. Such proximity suggests potential crosstalk between human T-cells and myeloid subsets within the tumor niche in humanized GBM PDOXs, in line with recent work showing that T-cell-myeloid partnerships can either sustain or suppress anti-GBM immunity^41^.

Together, introduced human CD45^+^ subsets can home to GBM tumors developing in the brain of humanized mice, assembling into structured immune clusters containing both lymphoid and myeloid populations. The predominance of CD4^+^ T-cells and CD11c^+^ myeloid cells, along with their close spatial association, reflects the establishment of organized immune niches that parallel those observed in human GBM.

### Human CD4^+^ T-cells infiltrating intracranial GBM tumors locally acquire features of exhausted, memory and regulatory T-cells

We further examined the differentiation, activation, and exhaustion status of tumor-infiltrating and spleen CD4^+^ T-cells in HU-CD34^+^ and HU-PBMC PDOX T188 (**Fig. 4A-D, S6A-E)**. We categorized CD4^+^ T-cells into two subsets: FoxP3^-^ conventional T-cells (Tconv) and FoxP3^+^ Tregs (**Fig. 4A, S6A**). Tconv cells predominantly exhibited PD-1^+^LAG-3^-^ and PD-1^-^LAG-3^-^ phenotypes (**Fig. 4B**). Tregs were primarily PD-1^-^LAG-3^-^, with a lower PD-1^+^LAG-3^-^ subset (**Fig. S6C**). Only a small fraction of CD4^+^ T-cells was PD-1^+/-^LAG-3^+^. Since the majority of Tconv and Treg cells in the spleen were PD-1^-^LAG-3^-^ (**Fig. 4B, S6C**), these data suggest the local phenotypic adaptation within the GBM tumor.

**Figure 4.**
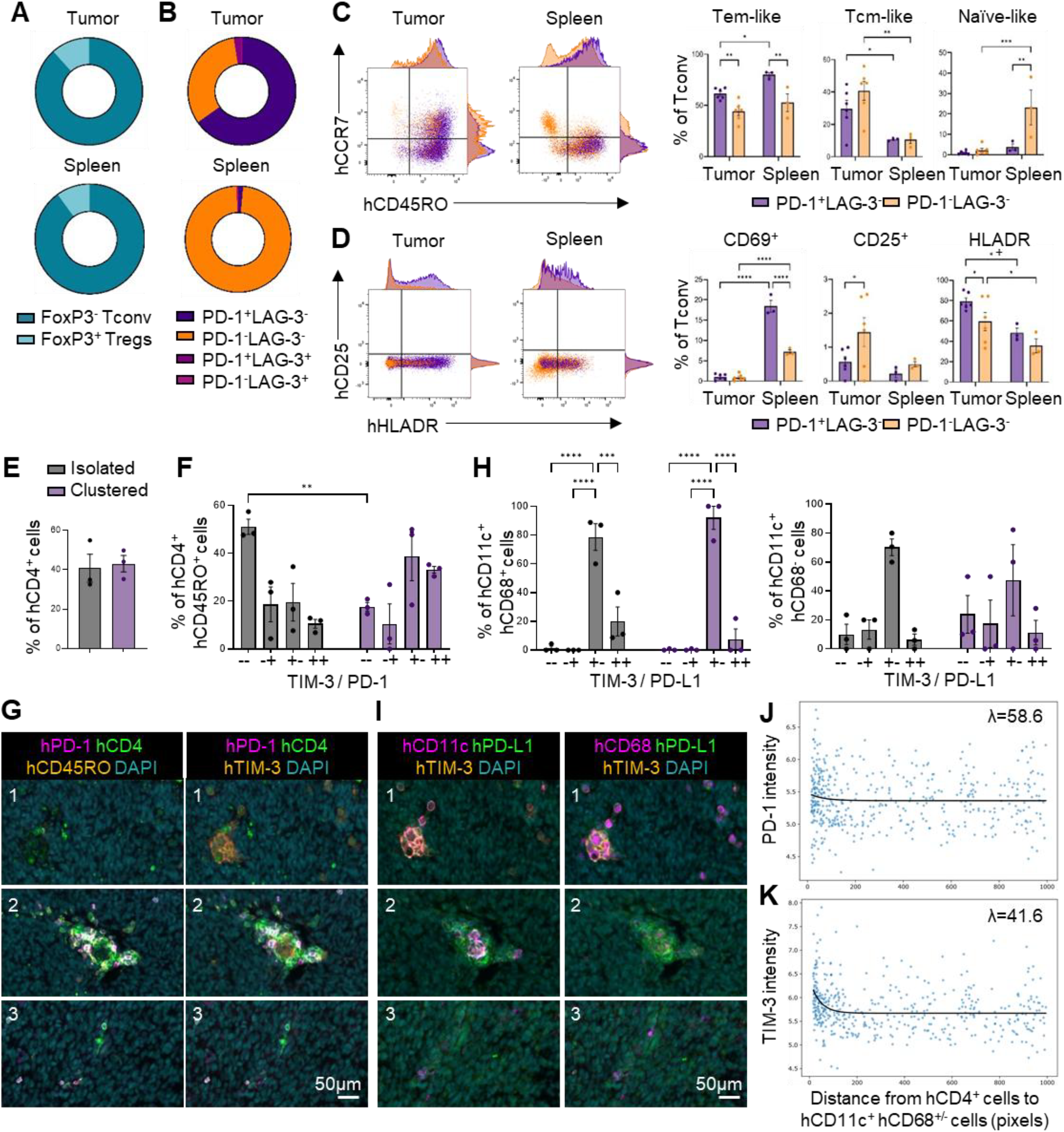
CD4^+^ T-cells and CD11c^+^ myeloid cells in humanized GBM PDOXs acquire immunosuppressive and spatially organized phenotypes within the TME. **A.** Composition of CD4^+^ T-cells categorized into FoxP3^-^ T conventional (Tconv) cells and FoxP3^+^ regulatory T-cells (Tregs) in HU-CD34^+^ PDOX T188 brain and spleen (n = 6 for brain, n = 3 for spleen, mean is shown). **B**. PD-1/LAG-3-based exhaustion states in Tconv cells in HU-CD34^+^ PDOX T188 (n=6 for tumor, n= 3for spleen, mean is shown). **C-D**. Representative flow cytometry dot plots and histograms display expression of (C) T-cell differentiation and (D) activation markers in PD-1^+^LAG-3^-^ and PD-1^-^LAG-3^-^ subsets in Tconv cells from tumor and spleen of HU-CD34^+^ PDOX T188. Quantification of (C) T-cell differentiation and (D) activation status in CD4^+^ Tconv cells from HU-CD34^+^ PDOX T188 brain tumors and spleens, PD-1^+^LAG-3^-^ exhausted and PD-1^-^LAG-3^-^ non-exhausted states are compared. T-cell differentiation classified central memory-like (Tcm; CD45RO^+^CCR7^+^), effector memory-like (Tem; CD45RO^+^CCR7^-^) and naïve-like (CD45RO^-^CCR7^+^) cells. T-cell activation was evaluated through CD69, CD25 and HLADR expression. For the gating strategy, see **Fig. S6A**. Statistical comparisons were made between PD-1^+^LAG-3^-^ and PD-^-^LAG-3^-^ subsets within tumors and spleen separately, as well as between tumor and spleen, two-way ANOVA (n=6 mice for PDOX tumor, n=3 for PDOX spleen, mean ± SEM, *p<0.05, **p<0.01, ***p<0.001, ****p<0.0001). **E**. Bar plot showing the distribution of CD45RO^−^ and CD45RO^+^ subsets among isolated and clustered human CD4^+^ T-cells. **F**. Distribution of TIM-3 and PD-1 expression within CD45RO^+^ CD4^+^ T-cells, comparing isolated and clustered populations using two-way ANOVA (n=3; mean ± SEM; ****p<0.0001). **G**. Representative multiplex immunofluorescence of the three ROIs shown in **Fig. 3E**, highlighting hPD-1 (pink), hCD4 (green), hCD45RO or hTIM-3 (orange), and DAPI (cyan). Scale bar: 50 µm. **H**. Distribution of TIM-3 and PD-1 expression within hCD11c^+^hCD68^+/-^ cells comparing isolated versus clustered using two-way ANOVA (n=3; mean ± SEM; ***p<0.001; ****p<0.0001). **I**. Representative multiplex immunofluorescence of the three ROIs as in **Fig. 3E**, showing hCD11c or CD68 (pink), hPD-L1 (green), hTIM-3 (orange), and DAPI (cyan), scale bar: 50 µm. **J-K**. Distance-intensity plots showing PD-1 (**J**) and TIM-3 (**K**) expression levels in hCD4^+^ T-cells relative to their distance from hCD11c^+^hCD68^+^/^−^ myeloid cells within Vimentin^+^ tumor regions. Each point represents a single CD4^+^ T-cell. Distances are shown in pixels. The black line indicates the overall exponential fit.

To trace CD4^+^ T-cell differentiation patterns, we distinguished putative central memory T-cells (Tcm-like, CD45RO^+^CD45RA^-^CCR7^+^), effector memory T-cells (Tem-like, CD45RO^+^CD45RA^-^CCR7^-^), effector memory CD45RA^+^ T-cells (Temra-like, CD45RO^-^ CD45RA^+^CCR7^-^) and naïve T-cells (Naïve-like, CD45RO^-^CD45RA^+^CCR7^+^, **Fig. S6A**). PD-1^+^LAG-3^-^ Tconv cells in tumors predominantly exhibited putative Tem-like and Tcm-like phenotypes, and high HLA-DR positivity (**Fig. 4C-D**). The rare PD-1^+^LAG-3^-^ Tconv cells in the spleen were largely in the Tem-like state. These data suggest potential activation of human T-cells in response to antigen encounter within the TME and/or peripheral T-cell priming. While HLA-DR indicates an active antigen response, PD-1 serves as an inhibitory receptor to prevent overactivation, which can lead to exhaustion. Their co-expression suggests that subsets of memory T-cells underwent a complex regulated activation program, potentially reflecting T-cell adaptation to the immunosuppressive signals of GBM and its local TME. Activation status was assessed based on CD69 (early), CD25 (intermediate), and HLA-DR (late) activation markers. PD-1^+^LAG-3^−^ Tconv cells exhibited a more activated HLA-DR^+^ phenotype in both tumor and spleen (**Fig. 4D**). In contrast, CD69^+^ cells were predominantly observed in the spleen, indicating early systemic activation signals less evident in the tumor. In contrast, PD-1^−^LAG-3^−^ Tconv cells displayed a higher proportion of naïve-like cells and reduced Tem-like and HLA-DR^+^ subsets (**Fig. 4C-D**), consistent with limited antigen exposure and activation. Both PD-1^+^LAG-3^-^ and PD-1^-^LAG-3^-^ Tregs showed similar differentiation states composed mainly of Tcm-like and Tem-like subsets with HLA-DR^+^ expression enriched in tumors compared to spleen (**Fig. S6D-E**). Overall, these data highlight the significant adaptation of CD4^+^ T-cells to the local TME, via a GBM-specific transition from naïve to effector/central memory phenotypes or a selective trafficking from circulation.

### Intratumoral human immune clusters create immunosuppressive niches enriched for exhausted CD4^+^ T-cells and immunosuppressive myeloid cells

We further examined the spatial distribution of CD4^+^ T-cells and myeloid subsets (**Fig. 4E-I**). Among clustered CD4^+^ cells, ∼40% expressed CD45RO (**Fig. 4E**) and the majority of these memory T-cells co-expressed TIM-3 and variably PD-1 (**Fig. 4F-G**), confirming the acquisition of exhaustion within the tumor milieu suggested by flow cytometry. Isolated CD4^+^ cells showed similar CD45RO^+^ positivity (∼40%, **Fig. 4E**), yet this subset had a significantly larger TIM-3^−^PD-1^−^ population (**Fig. 4F**), suggesting that clusters favor the acquisition of dysfunctional phenotypes.

Myeloid populations also displayed pronounced immunosuppressive features. Around 80% of CD11c^+^CD68^+^ cells expressed TIM-3 but lacked PD-L1 regardless of their spatial organization, while ∼20% co-expressed TIM-3 and PD-L1 (**Fig. 4H-I**). CD11c^+^CD68^−^ cells showed more heterogeneous patterns, still with predominance of the TIM-3^+^PD-L1^−^ subset. Together, these data reveal that human myeloid cells within GBM PDOXs acquire a heterogeneous yet predominantly immunosuppressive profile. While we cannot conclusively confirm subtype identity from multiplex imaging alone, the presence of CD11c^+^CD68^+/-^cells in close proximity to CD4^+^ T-cells was associated with stronger PD-1 and TIM-3 expression on the latter, suggesting the potential of TAM-like immunoregulatory phenotypes within the tumor (**Fig. 4J-K**). Overall, we show that human CD4^+^ T-cells in GBM PDOXs undergo localized phenotypic adaptation, acquiring memory-like features alongside signs of partial exhaustion. These shifts occur preferentially within immune clusters, where close spatial interaction with TIM-3^+^ myeloid cells may contribute to functional T-cell suppression. Collectively, these data demonstrate that humanized mouse models faithfully reproduce the hallmark features of the GBM immunosuppressive TME, defined by limited T-cell infiltration and dominant myeloid activity.

### Mouse-derived Mg acquires GBM-specific states in humanized PDOX mice

Since previous studies have highlighted complex interactions between T-cells and Mg in GBM tumors^41-43^, we explored whether the infiltration of human immune cells into GBM tumors in humanized PDOXs influenced mouse innate immune responses. Iba1^+^ Mg in HU-CD34^+^ control brains displayed a homeostatic, resting ramified morphology, which transitioned to an activated GBM-specific amoeboid morphology in the tumor-bearing HU-CD34^+^ PDOXs (**Fig. S6F**). This activation also occurred in close spatial proximity between Iba1^+^ mouse-derived TAMs and infiltrating hCD45^+^ immune cells in HU-CD34^+^ PDOX T188 (**Fig. S6G**). Mg activation towards CD45^high^ and CD11c^+^ TAMs was similar to PDOXs in NSG mice (**Fig. 2E, S6H**). An activation gradient was detected from control brains, to the distant regions and tumor areas similar to our previous profiling in Nude mice^20^. Still, neighborhood enrichment analysis (**Fig. 3G**) revealed that human CD4^+^ T-cells preferentially localized in proximity to human-derived myeloid cells (CD11c^+^CD68^+^/^−^) rather than to mouse Iba1^+^ cells. These findings confirm that humanized models can effectively support human immune responses without compromising murine immunity. An evaluation of the extent of the functional crosstalk between human lymphoid and mouse myeloid compartments will require further investigation.

### Targeting PD-1 and GITR modulates human immune cell infiltration to intracranial tumors

Given the limited clinical success of immune checkpoint blockade in GBM patients^11^, we aimed to evaluate whether humanized PDOXs can serve as a reliable platform to test immunotherapies with measurable modulation of human immune cells. As we detected intratumoral CD4^+^ T-cells, polarized towards Tconv and Treg phenotypes, we selected PD-1 and GITR as targets. This therapeutic combination is currently under clinical investigation in recurrent GBM patients (NCT04225039)^44^. *PDCD1* (PD-1) in human GBM patients (scRNA-seq, GBMap)^45^ is detected across several T-cell subsets, whereas *TNFRSF18* (GITR) is primarily restricted to Tregs and, to a lesser extent, NK cells (**Fig. S7A**). We confirmed expression of GITR on both Tconv and Treg cells in the tumor tissue of HU-CD34^+^ T188 PDOX, with enrichment in Tregs, while PD-1 was more prominent on Tconv cells (**Fig. S7B**).

We assessed four treatment arms: isotype control (IgG4κ + IgG1κ), pembrolizumab (anti-PD-1), ragifilimab (anti-GITR), and combination treatment (**Fig. 5A**). As anticipated from clinical trials^8-10^, neither therapy altered tumor growth kinetics compared to isotype-treated controls (**Fig. 5B-C**). Interestingly, we observed increased proportion of intratumoral hCD45^+^ cells upon anti-PD-1 treatment, but not anti-GITR (**Fig. 5D**). Infiltration was markedly lower in the contralateral (distant) hemisphere, suggesting that the immuno-modulatory effect was spatially restricted to the TME. The combination of anti-PD-1 and anti-GITR resulted in an intermediate phenotype, suggesting that addition of GITR may attenuate anti-PD-1-driven infiltration.

**Figure 5.**
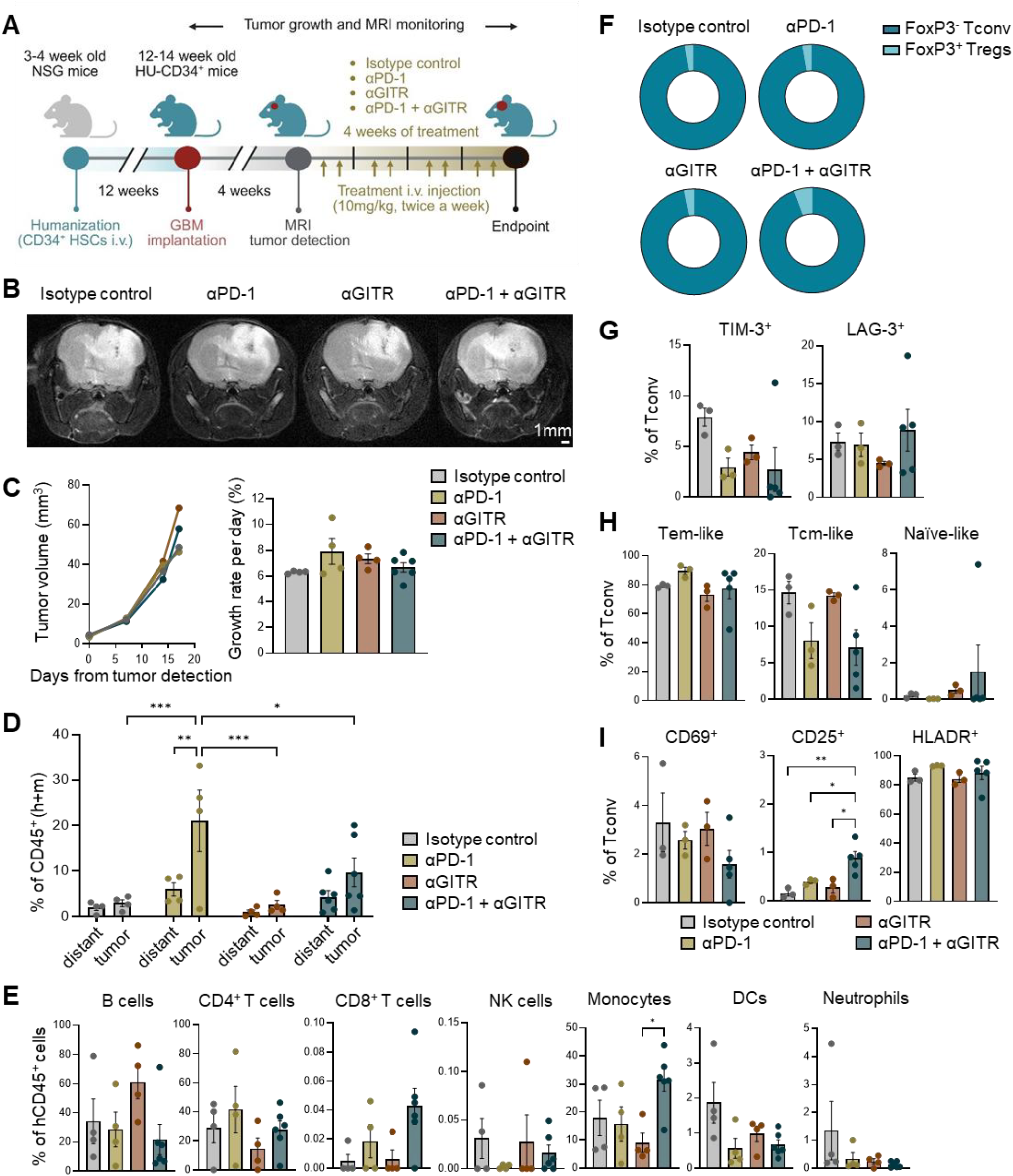
PD-1 and GITR targeting impact the immune landscape of humanized GBM PDOXs without altering tumor growth. **A.** Schematic illustrating treatment regimen in HU-CD34^+^ T188 PDOX mice. **B**. Representative T2-weighted MRI images of coronal brain sections from HU-CD34^+^ T188 PDOX mice across treatment arms, scale bar: 1 mm. **C**. Quantification of MRI-based tumor volume (mm^3^) and growth rate upon treatment. Left: each connected line represents the mean tumor volume per treatment group over time. Right: tumor growth rate was calculated between the time of first detection and experimental endpoint, one-way ANOVA with Tukey’s HSD correction (n=4 for isotype control, PD-1 and GITR monotherapy; n=6 for combination treatment; mean ± SEM; ns=not significant). **D**. Proportions of human (hCD45^+^) immune cells in the tumor-bearing (‘tumor’, right hemisphere) and distant (‘distant’, left hemisphere) brain regions across treatment arms, two-way ANOVA (n=4 for isotype control, PD-1 and GITR; n=6 for combination; mean ± SEM; *p<0.05, **p<0.01, ***p<0.001). **E**. Flow cytometry quantification of human lymphoid and myeloid subsets among human CD45^+^ cells in the tumors (right hemisphere) across the different treatment groups, one-way ANOVA with Tukey’s HSD correction (n=4 for isotype control, PD-1 and GITR; n=6 for combination; mean ± SEM; *p<0.05). See gating strategy in **Fig. S3C. F**. Proportions of CD4^+^ T-cells classified as FoxP3^-^ T conventional (Tconv) and FoxP3^+^ regulatory T-cells (Tregs) in HU-CD34^+^ PDOX T188 brains across treatment arms (n=3 for isotype control, PD-1 and GITR; n=5 for combination; mean shown for each group). **G-I**. Flow cytometry analysis of Tconv cells from HU-CD34^+^ PDOX T188 across treatment arms. (**G**) Quantification of exhaustion markers TIM-3 and LAG-3. (**H**) Distribution of differentiation subsets, defined as effector memory-like (Tem; CD45RO^+^CCR7^−^), central memory-like (Tcm; CD45RO^+^CCR7^+^), and naïve-like (CD45RO^−^CCR7^+^) cells. (**I**) Expression of activation markers CD69, CD25, and HLA-DR, one-way ANOVA with Tukey’s HSD correction (n=3 for isotype control, PD-1 and GITR; n=5 for combination; mean ± SEM; *p<0.05, **p<0.01).Gating strategy shown in **Fig. S6A**.

Human CD45^+^ cells in the tumor (**Fig. 5E**) and distant brain regions (**Fig. S7C**) presented only subtle treatment-dependent variations. We did not observe a selective infiltration of a specific cell type. B cells, CD4^+^ T-cells, and monocytes remained the dominant human immune populations, indicating a conserved immune cell profile irrespective of treatment. Only minor non-significant trends were observed: anti-PD-1-treated tumors showed increased CD4^+^ T-cells, GITR-treated tumors displayed increased B cells, whereas combination treatment enriched for monocytes (**Fig. 5E**), suggesting a subtle rebalance of lymphoid and myeloid composition. Murine Mg-TAMs were not affected, showing the same activation towards CD45^high^ and CD11c^+^ states in tumor-bearing hemispheres (**Fig. S7D**).

Systemic immune profiles were largely preserved, with only modest treatment-dependent trends. In bone marrow, blood and spleen, PD-1-based therapy favored higher CD4^+^ T-cells. GITR treatment was associated with increased B cells, and combination therapy showed a slight increase in monocytes (**Fig. S7E-G**). In cervical lymph nodes, control mice exhibited high monocyte proportions (**Fig. S7H**), consistent with prior reports showing that myeloid cells dominate cervical lymph nodes under steady-state conditions^46^ and play an important role in GBM immunoregulation^47^. In contrast, all treatment arms showed increased B and T-cells, particularly CD4^+^ T-cells in PD-1 and combination arms, likely reflecting enhanced priming and activation, potentially facilitating CD4^+^ T-cell recruitment into the brain. Overall, local and systemic immunity remained largely stable, yet subtle trends underscore the model’s ability to capture treatment-associated modulation relevant for translational immunotherapy studies. Cell type-unspecific increase of hCD45^+^ cells could be an indication of the higher BBB penetration within TME.

### Intratumoral CD4^+^ T-cells exhibit subtle phenotypic and functional shifts upon immunotherapy

We next examined the activation state of intratumoral CD4^+^ Tconv and Treg cells after treatment. The overall balance between Tconv and Tregs remained relatively stable across treatment arms (**Fig. 5F**). While epitope expression in isotype control mice was consistent with untreated HU-CD34^+^ GBM PDOXs, assessment of PD-1 and GITR expression in the treatment arms was technically limited, as therapeutic antibody binding masked detection of the epitopes (**Fig. S8A-B**).

Additional exhaustion markers TIM-3 and LAG-3 were detected on only small fractions of CD4^+^ T-cells, with combination therapy showing a trend towards reduced TIM-3 expression in Tconv and Tregs (**Fig. 5G, S8D**). This suggests that dual PD-1/GITR targeting may partially limit or reverse the acquisition of exhaustion features. Differentiation and activation profiles were largely preserved. Most Tconv and Tregs displayed Tem-like phenotypes, with only modest trends toward increased Tem-like Tconv proportions in PD-1-treated mice and reduced Tem-like Tregs in GITR-treated mice (**Fig. 5H, S8E**). Activation status was dominated by HLA-DR^+^ late-stage cells across all groups, with early and intermediate stages remaining scarce (**Fig. 5I, S8F**). Together, intratumoral CD4^+^ T-cells maintain stable activation and differentiation profiles after immunotherapy, with subtle trends in exhaustion and memory-associated states. These modest shifts are consistent with reported mechanisms of PD-1 and GITR modulation in other cancers^48-50^ and support the utility of HU-CD34^+^ GBM PDOXs as a platform to assess clinically relevant immunotherapies in a human immune context.

## Discussion

*In vivo* preclinical modelling is particularly challenging for solid tumors arising in hard-to-access organs, such as the brain. Developing patient-derived models that faithfully capture the complexity of tumor-immune interactions in GBM is essential for rational immunotherapy development. Although orthotopic xenografting offers replication of the organ-specific TME, efficient engraftment requires an immunodeficient background, which compromises a functional adaptive immunity and restricts their use for evaluating novel immunotherapies^51^. We have previously shown that PDOXs recreate a functional crosstalk where human GBM cells reciprocally interact with the mouse TME, including vasculature and innate immunity, undergoing significant molecular shifts toward GBM-specific states that can be modulated upon therapeutic interventions^18-22,27,52^. In this study, we further systematically characterized baseline and PDOX brain immunity across the most commonly used mouse strains, providing a reference framework for interpreting tumor-immune interactions. We demonstrated that GBM PDOXs established in humanized mice can recreate the hallmarks of human adaptive immunity within the brain. Both HU-CD34^+^ and HU-PBMC PDOXs supported systemic and local reconstitution of human immune cells, including intratumoral memory-like and partially exhausted CD4^+^ T-cells and a small subset of Tregs, closely resembling those in patient tumors^53,54^. These lymphocytes were modulated locally, and organized into spatially structured clusters together with human CD11c^+^ myeloid cells, mirroring immune niches recently described in human GBM^40,41^. These findings refine our understanding of human T-cell-myeloid interactions within the brain, demonstrating how adaptive exhaustion also emerges under species-specific myeloid pressure. Functionally, immunotherapy mildly modulated human immune infiltration and states without altering murine TAM immunosuppression, aligning with the clinical reality of limited tumor control from single-agent checkpoint agents in GBM^11^ and underscoring the value of these models for translational immunotherapy testing. Together, these results confirm that humanized GBM PDOXs recapitulate key features of the GBM immune landscape, providing a meaningful framework to interrogate tumor-immune crosstalk and therapeutic modulation *in vivo*.

To our knowledge, this represents the first side-by-side comparison of HU-CD34^+^ and HU-PBMC systems, revealing model-specific advantages. Both models allow for introducing the adaptive immune system from CD34^+^ hematopoietic stem cells or PBMCs, respectively^31,51,55^. Such humanized mice have proven useful in interrogating autoimmunity^12,13^, and human infectious diseases^14,15^. In our hands, HU-CD34^+^ mice achieved higher and more durable levels of systemic humanization, with restoration of splenic white pulp and sustained hematopoietic activity in the bone marrow, while HU-PBMC mice yielded rapid lymphoid reconstitution with limited myeloid output. Both systems supported intracranial tumor engraftment, confirming that the human immune compartment does not prevent GBM outgrowth, in agreement with previous humanized GBM reports using hu-BLT or hu-NOG-EXL hosts^56-58^. Notably, the immune composition in our models converged with human GBM, marked by the selective attraction of CD4^+^ T-cells, myeloid and B cells, highlighting the reproducibility of the GBM-associated immune landscape even under distinct modes of humanization. Although human CD8^+^ T and NK cells were present systemically, they did not significantly infiltrate intracranial tumors, aligning with clinical data which shows GBM is largely resistant to cytotoxic immune responses^59^. The observation that murine TAMs remain dominant further reinforces the conserved myeloid dependency of GBM growth^2,20,41,42^.

Our comparative approach also clarifies each humanization strategy’s experimental scope. HU-CD34^+^ mice allow long-term studies of systemic and local immunity, including bone marrow and lymph node interactions. HU-PBMC mice, despite GvHD, remain useful for short-term immunotherapy assessments. The limited infiltration of human monocytes and CD8^+^ T-cells observed highlights a current challenge shared across humanized models^55^. Future improvements, including transgenic expression of human cytokines (e.g., IL-3, GM-CSF, SCF)^60,61^, adoptive supplementation with iPSC-derived myeloid cells^62^, or the use of HLA-transgenic NSG backgrounds^63,64^ may enhance myeloid and antigen-presenting cell development, thereby increasing the accuracy of antigen-specific immune responses. Nonetheless, the present models already reproduce key features of human GBM immunity, including T-cell exhaustion, regulatory T-cell enrichment, and dominant myeloid suppression, reflecting the true immune landscape of patient tumors ^3,4,53^. Despite higher cost, they offer a more physiologically relevant context for immunotherapy testing over murine counterparts, given the fundamental differences between murine and human immune environments.

The responsiveness of the reconstructed immune system to immunotherapy strengthens its translational value. Treatment with anti-PD-1 or anti–GITR antibodies, alone or in combination, only mildly modulated human immune infiltration and phenotypic states within tumors without affecting the murine myeloid compartment. Interestingly, PD-1 blockade increased human CD45^+^ infiltration into GBM tumors, whereas anti-GITR alone did not and dual therapy produced an intermediate phenotype, suggesting distinct modulatory effects on the adaptive compartment. These changes were spatially restricted to the tumor areas and did not translate into growth control, in line with negative GBM trials^8-10^. In syngeneic mouse GBM models, dual PD-1/GITR targeting reprogrammed Tregs toward effector states and enhanced anti-tumor immunity^50^. Our humanized PDOX findings clearly differ, showing no growth benefit and only modest immune-state shifts, likely reflecting species-specific myeloid composition, distinct GITR signaling within the human T-cell compartment, and limited intratumoral CD8^+^ support compared to fully immunocompetent hosts, all of which represent biologically faithful constraints rather than simple model limitations. The ability of our system to mirror these therapeutic bounda ries emphasizes its suitability for dissecting context-dependent mechanisms of immunotherapy resistance and for evaluating rational combinations that prime the TME before checkpoint engagement^11^.

Interestingly, systemic immune composition remained largely stable upon treatment, yet the cervical lymph nodes exhibited the most consistent trends toward lymphoid enrichment, particularly under PD-1-based therapy. This observation aligns with recent work demonstrating that cervical nodes and skull bone marrow niches act as critical hubs for brain-immune communication and T-cell priming^65,66^. The capacity to capture therapy-related changes in these compartments points to future applications in exploring how lymphatic drainage and skull bone marrow-derived cells shape immune surveillance and treatment responsiveness in GBM. Similarly, the presence of human B and T-cells within tumors suggests the potential emergence of tertiary lymphoid-like structures (TLS), a feature recently reported in human gliomas^67,68^. Further investigation of TLS maturation and function in this system could clarify whether they act as immune-activating niches or rather contribute to tumor tolerance.

Overall, our findings place humanized GBM PDOXs as a next-generation preclinical platform that transcends conventional modeling by integrating patient-derived tumor biology with a functional human immune system in its native brain context. Beyond modeling, these systems provide mechanistic insights into the spatial and functional interplay between adaptive and innate immunity in GBM, revealing how they co-evolve to enforce tumor tolerance. Their application extends to other solid tumors characterized by immune exclusion and myeloid-driven suppression, including pancreatic, liver, and metastatic brain cancers^69-71^. The ability to read out human T-cell states in the correct organ context while preserving tissue-native vasculature and myeloid ecosystems enables realistic testing of combination immunotherapies, bispecifics, and cell therapies, including regional delivery proven feasible in patients^72,73^, with physiologic constraints that simpler systems miss.

In summary, humanized GBM PDOXs reproduce the core immune biology of human GBM: CD4-biased, memory/exhaustion-prone T-cells under immunosuppressive pressure, and respond to clinically relevant immune perturbations in ways that expose the true barriers to efficacy. By integrating tumor fidelity, human adaptive immunity, and organ-specific barriers they conform a versatile platform that can accelerate rational design and triage of next-generation immunotherapies and inform strategies for other difficult-to-treat solid tumors.

### Limitations of the study

This study’s limitations provide avenues for future improvement and refinement. While humanized GBM PDOXs successfully recreate key aspects of human adaptive immunity, several factors deserve consideration. Direct demonstration of antigen-driven T-cell activation was not possible and impact of mouse MHC on antigen presentation and T-cell education requires further attention. It remains to be seen if replacement of brain-resident Mg with human derivatives will influence the GBM immunity in humanized mice. While some mouse and human cytokines exhibit species cross-reactivity^74,75^, the specific interplay of human and mouse cytokines in humanized GBM models remains poorly understood. Nonetheless, the predominance of Tem- and Tcm-like phenotypes strongly suggests antigen encounter within the GBM microenvironment. This is consistent with the intrinsically low neoantigen load of GBM and the use of CD34^+^ HSC-based humanization, which while limiting measurable specificity, also recapitulates the biological reality of patient tumors. The use of targeted flow cytometry panels provided a reliable overview of key populations but likely underestimated minor suppressive subsets such as MDSCs. Finally, the relatively low human monocyte and CD8^+^ T-cell infiltration may restrict some immune interactions, yet these features mirror physiological constraints of GBM rather than technical artifacts. Overall, these caveats delineate the model’s physiological boundaries without undermining its translational relevance and highlight specific avenues for refinement in future studies.

## Acknowledgements

We thank the LIH PRECISION-PDX Brain Tumor Bank (www.precision-pdx.lu), supported by the patients, the Neurosurgery Department of the Centre Hospitalier de Luxembourg, the NORLUX team, and the Clinical and Epidemiological Investigation Center of LIH. We appreciate the support from LIH’s core facilities, including the Animal Facility, In Vivo Imaging Facility, and the National Cytometry Platform (NCP). We are grateful to Dr. Guy Berchem for providing Keytruda (pembrolizumab), Dr. Magretta Adiamah for proofreading and Hamed Allahverdi for bioinformatics support.

## Author contributions

Conceptualization: PMS, AO, SPN, AG; Methodology: PMS, AO, BK, HD, EK, VB, CR, CSD, MR, AP, AM, JP, EM, JHMP, SPN, AG; Investigation: PMS, AO, BK, HD, EK, VB, MR, AM, AG; Formal analysis: PMS, AO, BK, HD, AG; Resources: AO, CSD, JP, EM, AM, JHMP, SPN, AG; Supervision: AO, CSD, SPN, JHMP, AG; Writing - Original Draft: PMS, AO, AG; Writing - Review & Editing: all authors.

## Data availability

Molecular information on PDO/PDOX models is available via the PDMC Finder (https://www.cancermodels.org/about/providers/lih).

## Materials and Methods

### Clinical samples and PDOX cohort

GBM patients provided informed consent, and tumor collection procedures were approved by the local research ethics committees (National Committee for Ethics in Research (CNER) Luxembourg: 201201/06; Haukeland University Hospital: 2009/117). The establishment of the cohort of primary patient-derived organoids and PDOXs, as well as associated animal experimentation protocols have been previously described^18,19^. Briefly, primary organoids were generated from GBM tissue fragments for up to two weeks at 37°C in a 5% CO_2_ environment, using DMEM medium supplemented with 10% FBS, 2 mM L-Glutamine, 0.4 mM NEAA, and 100 U/ml Pen–Strep (all from Lonza). To establish PDOX models, GBM organoids were intracranially implanted following our standard protocol^19^. For regular PDOX maintenance, we implanted six organoids into the right frontal cortex of NSG (NOD.Cg-Prkdc^scid^ Il2rg^tm1Wjll^/SzJ) mice. All animal experiments were conducted following the regulations of the European Directive on animal experimentation (2010/63/EU) and approved by the Luxembourg Institute of Health’s Animal Welfare Structure and the Luxembourg Ministry of Agriculture (protocols LUPA2019/93, LUPA2024/10 and LUPA2019/67). Animals were housed in a specific pathogen free (SPF) facility with controlled temperature, humidity, and light conditions, and had water and food ad libitum. Five GBM PDOX models were used in this study: P3, P8, P13, T158 and T188 (**Table S1**)^18^.

### Animal experimentation in immunodeficient mouse strains

### Mouse strains

Three immunodeficient mouse strains were utilized. From lower to higher degree of immunodeficiency: Nude mice (Crl:NU(Ico)-Foxn1^nu^; absent mature T-cells), Rag1 KO mice (B6.129S7-Rag1^tm1Mom^/J; absent mature T and B cells) and NSG mice (NOD.Cg-Prkdc^scid^ Il2rg^tm1Wjll^/SzJ; absent mature T, B and NK cells) mice. C75BL/6J or C57BL/6N mice served as fully immunocompetent controls (**Fig. 1A**). All mice were obtained from Charles River Laboratories.

#### PDOX derivation and monitoring

PDOXs were established by intracranial implantation of GBM organoids in the right hemisphere of 12-week-old female mice (minimum 3 mice per model) from Nude, Rag1KO, and NSG strains^19^. Mice underwent weekly monitoring via T2-weighted MRI, and tumor volume quantified using ImageJ software. Mice were sacrificed upon reaching humane endpoint (tumor volume of 100 mm^3^ or loss of ≥15% baseline body weight)^19,76^. Mice were intracardially perfused with 10 mL of ice-cold PBS, and brains were collected for further assays.

### Orthotopic xenografting in humanized mouse models

#### HLA-typing

Human GBM cells of selected PDOX models were HLA typed with the TruSight™ v.2 HLA Sequencing Panel (Illumina) for ultrahigh resolution sequencing of 11 HLA loci (Class I HLA-A, -B, -C; Class II HLA-DPA1, -DPB1, -DQA, -DRB1/3/4/5) (**Table S1**). HLA type of immune cells was provided by the commercial vendors (**Table S2**).

#### Commercially sourced HU-CD34^+^ mice (gamma-ray protocol)

We purchased 10 HU-CD34^+^ female mice (12-weeks-old) from the Jackson Laboratory (JAX). These mice received 140cGY of gamma irradiation and were subsequently engrafted with 6*10^4^ CD34^+^ immune cells at 3-4 weeks of age. To minimize the risk of allogeneic responses, the CD34^+^ cell donor was partially HLA-matched with the PDOX T188 model used for tumor implantation (**Table S2**). The level of humanization was verified at 12 weeks (JAX) and 19 weeks (in-house) prior to implantation. HU-CD34^+^ mice were intracranially implanted with six T188 GBM organoids in the right hemisphere (n=7), while the remaining HU-CD34^+^ mice served as healthy controls (n=3).

#### In-house generated HU-CD34^+^ mice (Busulfan protocol)

HU-CD34^+^ mice were generated in-house based on adapted protocol from Adams et al.^77^. Briefly, 3-4 week-old female NSG mice were treated with two doses of 20 mg/kg of busulfan (Merck) 48h and 24h before injection with 5-10*10^4^ CD34^+^ commercial human cord blood cells (Lonza; animal protocol LUPA2019/67, no HLA typing available, **Table S2**) via the lateral tail vein. After 10-12 weeks, the humanization level was checked in peripheral blood, and mice were considered humanized if showing >25% human CD45^+^ immune cells in the circulation. 12-14 week-old mice were intracranially implanted with T188 or T158 GBM organoids^18^ (n=3 per PDOX for characterization, n=18 for T188 PDOX treatment study).

#### In-house generated HU-PBMC mice

12-week-old NSG female mice were intracranially implanted with T188 or T158 GBM organoids^18^. Upon detection of tumor growth by MRI, peripheral blood mononuclear cells (PBMCs, Immunospot, partially matched to T188 and T158, **Table S2**) were injected at 10^7^ cells per 100 μL of RPMI into the lateral tail vein of GBM PDOXs or healthy NSG counterparts (n=3 in control groups and n=7 in implanted groups).

#### Monitoring and sample collection for humanized GBM PDOXs

All humanized mice were monitored weekly by MRI and checked for weight loss. Mice were sacrificed at the endpoint defined by tumor growth progression or, in the case of HU-PBMC mice, upon presentation of graft-versus-host disease (GvHD) symptoms (i.e., weight loss, hair loss, anemia, or gastrointestinal signs^30,76^). All mice were approximately 22-25 weeks old at the study endpoint, when animals were perfused with ice-cold PBS, and brains were collected along with spleen, bone marrow, and blood.

### Immunotherapy in humanized GBM PDOXs

T188 GBM organoids were implanted in HU-CD34^+^ mice after 12 weeks of humanization and tumor growth was confirmed 4 weeks later by MRI. 18 HU-CD34^+^ PDOX T188 mice were randomly assigned to four treatment arms based on MRI tumor detection: isotype control (human IgG1κ and IgG4κ, MedChemExpress; n=4), anti-human PD-1 monotherapy (pembrolizumab, hIgG4, MSD; n=4), anti-human GITR monotherapy (ragifilimab, hIgG1, MedChemExpress; n=4), or a combination of anti-human PD-1 and GITR (n=6). All treatments were administered intravenously at a dose of 10 mg/kg, twice per week, for 4 consecutive weeks. At endpoint, mice were euthanized and perfused with ice-cold PBS. Brain, spleen, bone marrow, cervical lymph nodes, and blood were collected.

### Magnetic resonance imaging (MRI)

Anesthesia was induced and maintained with a 2.5% isoflurane/oxygen mixture (CP Pharma). Mice were positioned in a 3T MRI system (MRsolution) with continuous monitoring of respiration and body temperature. Initial imaging used a T2-weighted scan (which highlights fluid-filled regions and tumors) for multi-slice axial acquisition. Tumor volume was quantified from hyperintense regions in T2-weighted axial MRI slices measured using ImageJ. Total tumor volume was obtained in mm^3^ by adding up the tumor areas across each slice (1 mm thickness each), following our established protocols^19,76^. The daily growth rate was calculated as the percentage change in tumor volume (mm^3^) per day based on MRI, determined by the logarithmic difference between the final tumor volume and the initial tumor volume, divided by the total number of days.

### Multicolor flow cytometry

#### Tissue and blood processing

PDOX tumor tissue was dissected according to MRI and histopathological features. Left and right hemispheres were separated, when specified. Tumor and control brain tissues were dissociated with the MACS Neural Tissue Dissociation Kit (P) (Miltenyi Biotec), following manufacturer’s protocol. Single cells were purified using the Myelin Removal Beads II kit (Miltenyi Biotec). For PDOXs in immunodeficient mice, human GBM cells were separated from the mouse TME using the Mouse Cell Depletion kit (Miltenyi Biotec) as previously described^19^. Spleens and lymph nodes from humanized GBM PDOXs were processed on a petri dish moistened with RPMI without supplements. Tissue was gently minced on a 70 µm cell strainer using the flat end of a 5 mL syringe plunger. Single-cell suspension was filtered with 50µm CellTrics (Sysmex) and washed with RPMI. Bone marrow was collected from mouse femurs by inserting a 26G needle and flushing with 10 mL of ice-cold RPMI into a 50 mL Falcon tube. The bone was discarded after flushing, and cells were resuspended, filtered with 50µm CellTrics (Sysmex). Cells were washed once with RPMI before downstream applications or cryopreservation. Approximately 250 µL of blood per mouse was collected via the submandibular vein into EDTA-coated tubes to prevent clotting. For flow cytometry, 50 µL of blood was placed in FACS tubes. Remaining blood was centrifuged at 3000 rpm for 10 min at 4°C to separate plasma, which was stored at -80°C for later analysis.

#### Flow cytometry-based acquisition

Antibodies applied are listed in **Table S3**. Single-cell suspensions were prepared in ice-cold FACS buffer (1X PBS + 0.2% BSA, 100 µl/test). Fc receptors were blocked using (i) anti-mouse CD16/CD32 antibody (Biolegend) for 10 min at 4°C, or (ii) Human TruStain FcX (Biolegend) for 10 min at room temperature. Cells were incubated with pre-conjugated antibodies for 30 min at 4°C in the dark. Bone marrow and blood samples were treated 5 min on ice or 15 min at RT, respectively, with Red Blood Cell Lysis buffer (Biolegend). For hFoxP3 staining, samples were permeabilized and fixed with True-Nuclear™ Transcription Factor Buffer Set (Biolegend). Data acquisition was performed on a NovoCyte Quanteon flow cytometer (Agilent) equipped with 405nm (violet), 488nm (blue), 561nm (yellow-green), and 640nm (red) lasers or on a BD FACSymphony S6 Cell Sorter (BD), with 355nm (ultraviolet), 405nm (violet), 488nm (red), 561nm, (yellow-green) and 637nm (red) lasers. Data was analyzed using FlowJo software (version 10.9.0) and visualized in GraphPad Prism 10. Each biological replicate represents an individual mouse.

### *Ex vivo* T-cell stimulation with PMA/Ionomycin

Human CD3^+^ T-cells were isolated from the spleens of humanized mice using a two-step protocol involving the Mouse Cell Depletion Kit and Pan T-cell Isolation Kit (both Miltenyi). CD3^+^ T-cells were seeded into 96-well U-bottom plates (Greiner) in RPMI GlutaMAX medium (ThermoFisher), supplemented with 9% FBS (Sigma), 1% human serum (Sigma), 0.1 mM sodium pyruvate (ThermoFisher), 5 mM HEPES (ThermoFisher), 55 µ M β-mercaptoethanol, 0.4 mM NEAA (Lonza), and 100 U/mL Pen–Strep (Lonza). Cells were incubated for 1h at 37°C in a 5% CO_2_ environment and stimulated with 50 ng/µL PMA and 1 µg/mL ionomycin (both Sigma) per well. To assess T-cell degranulation, CD107a antibody (BioLegend, **Table S3**) was added simultaneously with PMA/Ionomycin. After a 1h incubation, GolgiStop (1:1000) and GolgiPlug (1:500) (both BD) were added to the wells, followed by a 4h incubation at 37°C. The cells were then washed and stained for flow cytometric analysis using a specific cytokine panel (**Table S3**).

### Immunohistochemistry

Tumor-bearing PDOX brains were stained with H&E and GBM-specific markers Nestin and Vimentin, following previously established protocols^18,27^. Antibodies are listed in **Table S3**. Formalin-fixed paraffin-embedded (FFPE) brains were cut with a microtome into 3-4 µm coronal sections. Slide sections were heated at 95°C for 30 min in antigen retrieval solution (Dako) followed by primary antibody incubation either overnight at 4°C or for 3h at room temperature. This was followed by secondary antibody incubation with signal development using the Envision+ System/HRP Kit over 5–20 min (K4007, Agilent/Dako) for 3 min. Hematoxylin was used as a nuclear counterstain to enable visualization of tumor regions. Imaging was performed on a Nikon Ni-E microscope, capturing multiple fields of the tumor core.

### CellDive multiplex immunofluorescence

3-4 µm thick FFPE brain sections were mounted onto Superfrost™Plus slides. Antigen retrieval was performed on the Bond RXm system (Leica Biosystems, UK) as described^78^. Slides were stained with DAPI, mounted in Clickwells, and imaged using the CellDive platform (Leica Microsystems, Germany) equipped with a BAB 200 robot (Advanced Solutions, USA). Imaging was conducted at 2× (overview), 10× (region selection), and 20× (high resolution) magnifications. After acquiring autofluorescence and DAPI, slides underwent sequential antibody staining and imaging cycles, with background subtraction and fluorescence quenching (20% 0.5 M NaHCO_3_ + 10% H_2_O_2_ in ddH_2_O) performed twice per cycle^78,79^. From the second round onward, only conjugated antibodies were used (**Table S3**). DAPI served to stitch and align tiled images. Fluorescence detection employed filters optimized for DAPI, Alexa Fluor 488, 555/Cy3, 647, and 750 channels, using a penta-bandpass polychroic mirror and single-bandpass emission filters matched to the illumination source (**Table S4**). Single-cell segmentation and feature extraction were performed with HALO AI (IndicaLabs). Marker positivity was determined using global and local thresholding in Python (*scikit-image*^80^). For each antibody channel, global thresholds were set via Otsu’s method^81^ or mean + k·SD, and verified visually. Cells were considered positive if their mean intensity exceeded threshold values and cytoplasmic completeness met minimal cutoffs (10–15% for immune markers; ≥60% for IBA1, Vimentin, SMA). For mutually exclusive phenotypes, weighted scores combining intensity and staining completeness resolved assignments. To correct for regional artefacts, local adaptive thresholds^82^ were applied, requiring signal to exceed both global and local cutoffs. Local threshold maps were generated for quality control. Spatial neighbor graphs and enrichment analyses were computed in Python using Squidpy library^83^. For spatial proximity analyses, λ value shows the empirical range of influence.

### Statistical analyses

Statistical analyses and sample sizes (n) are provided in the figure legends. For comparisons between two groups, an unpaired, two-tailed Student’s t-test was used; to compare the proportions a Bonferroni multiple-significance-test correction was applied. When assessing more than two groups, we applied a one-way analysis of variance (ANOVA) with Tukey’s Honest Significant Difference (HSD) correction for multiple comparisons. For experiments involving comparing impact of two independent variables, a two-way ANOVA was used to assess interaction effects. Data points represent biological replicates, shown as individual dots, with error bars indicating standard error mean (SEM).

## References

1 Yabo, Y. A., Niclou, S. P. & Golebiewska, A. Cancer cell heterogeneity and plasticity: A paradigm shift in glioblastoma. Neuro Oncol 24, 669–682 (2022). 10.1093/neuonc/noab269

2 Sharma, P., Aaroe, A., Liang, J. & Puduvalli, V. K. Tumor microenvironment in glioblastoma: Current and emerging concepts. Neurooncol Adv 5, vdad009 (2023). 10.1093/noajnl/vdad009

3 Ravi, V. M. et al. T-cell dysfunction in the glioblastoma microenvironment is mediated by myeloid cells releasing interleukin-10. Nat Commun 13, 925 (2022). 10.1038/s41467-022-28523-1

4 Mohme, M. et al. Immunophenotyping of Newly Diagnosed and Recurrent Glioblastoma Defines Distinct Immune Exhaustion Profiles in Peripheral and Tumor-infiltrating Lymphocytes. Clin Cancer Res 24, 4187–4200 (2018). 10.1158/1078-0432.Ccr-17-2617

5 Chulpanova, D. S., Kitaeva, K. V., Rutland, C. S., Rizvanov, A. A. & Solovyeva, V. V. Mouse Tumor Models for Advanced Cancer Immunotherapy. Int J Mol Sci 21 (2020). 10.3390/ijms21114118

6 Zanella, E. R., Grassi, E. & Trusolino, L. Towards precision oncology with patient-derived xenografts. Nat Rev Clin Oncol 19, 719–732 (2022). 10.1038/s41571-022-00682-6

7 Connor, K., Golebiewska, A. & Byrne, A. T. Challenging the status quo to improve the translational potential of preclinical oncology studies. Nat Rev Cancer (2024). 10.1038/s41568-024-00756-w

8 Reardon, D. A. et al. Effect of Nivolumab vs Bevacizumab in Patients With Recurrent Glioblastoma: The CheckMate 143 Phase 3 Randomized Clinical Trial. JAMA Oncol 6, 1003–1010 (2020). 10.1001/jamaoncol.2020.1024

9 Omuro, A. et al. Radiotherapy combined with nivolumab or temozolomide for newly diagnosed glioblastoma with unmethylated MGMT promoter: An international randomized phase III trial. Neuro Oncol 25, 123–134 (2023). 10.1093/neuonc/noac099

10 Lim, M. et al. Phase III trial of chemoradiotherapy with temozolomide plus nivolumab or placebo for newly diagnosed glioblastoma with methylated MGMT promoter. Neuro Oncol 24, 1935–1949 (2022). 10.1093/neuonc/noac116

11 Moreno-Sanchez, P. M., Rezaeipour, M., Marit-Inderberg, E., Platten, M. & Golebiewska, A. Immunosuppressive mechanisms and therapeutic interventions shaping glioblastoma immunity. Nat Cancer in press (2025).

12 Kuwana, Y. et al. Epstein-Barr virus induces erosive arthritis in humanized mice. PloS one 6, e26630 (2011). 10.1371/journal.pone.0026630

13 Andrade, D. et al. Engraftment of peripheral blood mononuclear cells from systemic lupus erythematosus and antiphospholipid syndrome patient donors into BALB-RAG-2−/− IL-2Rγ−/− mice: A promising model for studying human disease. Arthritis & Rheumatism 63, 2764–2773 (2011). 10.1002/art.30424

14 Libby, S. J. et al. Humanized nonobese diabetic-scid IL2rγnull mice are susceptible to lethal Salmonella Typhi infection. Proceedings of the National Academy of Sciences 107, 15589–15594 (2010). 10.1073/pnas.1005566107

15 Sato, K. et al. A novel animal model of Epstein-Barr virus–associated hemophagocytic lymphohistiocytosis in humanized mice. Blood, The Journal of the American Society of Hematology 117, 5663–5673 (2011). 10.1182/blood-2010-09-305979

16 Berges, B. K., Akkina, S. R., Folkvord, J. M., Connick, E. & Akkina, R. Mucosal transmission of R5 and X4 tropic HIV-1 via vaginal and rectal routes in humanized Rag2−/− γc−/−(RAG-hu) mice. Virology 373, 342–351 (2008). 10.1016/j.virol.2007.11.020

17 Satheesan, S. et al. HIV replication and latency in a humanized NSG mouse model during suppressive oral combinational antiretroviral therapy. Journal of virology 92, 10.1128/jvi.02118-02117 (2018). 10.1128/jvi.02118-17

18 Golebiewska, A. et al. Patient-derived organoids and orthotopic xenografts of primary and recurrent gliomas represent relevant patient avatars for precision oncology. Acta Neuropathol 140, 919–949 (2020). 10.1007/s00401-020-02226-7

19 Oudin, A. et al. Protocol for derivation of organoids and patient-derived orthotopic xenografts from glioma patient tumors. STAR Protoc 2, 100534 (2021). 10.1016/j.xpro.2021.100534

20 Yabo, Y. A. et al. Glioblastoma-instructed microglia transition to heterogeneous phenotypic states with phagocytic and dendritic cell-like features in patient tumors and patient-derived orthotopic xenografts. Genome Medicine 16, 51 (2024). 10.1186/s13073-024-01321-8

21 Oudin, A., Moreno-Sanchez, P. M., Baus, V., Niclou, S. P. & Golebiewska, A. Magnetic resonance imaging-guided intracranial resection of glioblastoma tumors in patient-derived orthotopic xenografts leads to clinically relevant tumor recurrence. BMC Cancer 24, 3 (2024). 10.1186/s12885-023-11774-6

22 Fack, F. et al. Bevacizumab treatment induces metabolic adaptation toward anaerobic metabolism in glioblastomas. Acta Neuropathol 129, 115–131 (2015). 10.1007/s00401-014-1352-5

23 Shultz, L. D. et al. Human lymphoid and myeloid cell development in NOD/LtSz-scid IL2R gamma null mice engrafted with mobilized human hemopoietic stem cells. J Immunol 174, 6477–6489 (2005). 10.4049/jimmunol.174.10.6477

24 Flanagan, S. P. ‘Nude’, a new hairless gene with pleiotropic effects in the mouse. Genet Res 8, 295–309 (1966). 10.1017/s0016672300010168

25 Mombaerts, P. et al. RAG-1-deficient mice have no mature B and T lymphocytes. Cell 68, 869–877 (1992). 10.1016/0092-8674(92)90030-g

26 Nimmerjahn, A., Kirchhoff, F. & Helmchen, F. Resting microglial cells are highly dynamic surveillants of brain parenchyma in vivo. Science 308, 1314–1318 (2005). 10.1126/science.1110647

27 Bougnaud, S. et al. Molecular crosstalk between tumour and brain parenchyma instructs histopathological features in glioblastoma. Oncotarget 7, 31955–31971 (2016). 10.18632/oncotarget.7454

28 Benakis, C. et al. T cells modulate the microglial response to brain ischemia. Elife 11 (2022). 10.7554/eLife.82031

29 Klein, E., Hau, A. C., Oudin, A., Golebiewska, A. & Niclou, S. P. Glioblastoma Organoids: Pre-Clinical Applications and Challenges in the Context of Immunotherapy. Front Oncol 10, 604121 (2020). 10.3389/fonc.2020.604121

30 Elhage, A., Sligar, C., Cuthbertson, P., Watson, D. & Sluyter, R. Insights into mechanisms of graft-versus-host disease through humanised mouse models. Biosci Rep 42 (2022). 10.1042/bsr20211986

31 De La Rochere, P. et al. Humanized Mice for the Study of Immuno-Oncology. Trends Immunol 39, 748–763 (2018). 10.1016/j.it.2018.07.001

32 Huey, D. D. et al. Role of Wild-type and Recombinant Human T-cell Leukemia Viruses in Lymphoproliferative Disease in Humanized NSG Mice. Comp Med 68, 4–14 (2018).

33 Ishikawa, F. et al. Development of functional human blood and immune systems in NOD/SCID/IL2 receptor {gamma} chain(null) mice. Blood 106, 1565–1573 (2005). 10.1182/blood-2005-02-0516

34 Matas-Céspedes, A. et al. Use of human splenocytes in an innovative humanised mouse model for prediction of immunotherapy-induced cytokine release syndrome. Clin Transl Immunology 9, e1202 (2020). 10.1002/cti2.1202

35 Hess, N. J. et al. Inflammatory CD4/CD8 double-positive human T cells arise from reactive CD8 T cells and are sufficient to mediate GVHD pathology. Sci Adv 9, eadf0567 (2023). 10.1126/sciadv.adf0567

36 Adams, R. C. et al. Donor bone marrow–derived macrophage MHC II drives neuroinflammation and altered behavior during chronic GVHD in mice. Blood 139, 1389–1408 (2022). 10.1182/blood.2021011671

37 Lambert, N. et al. Central nervous system manifestations in acute and chronic graft-versus-host disease. Brain (2024). 10.1093/brain/awae340

38 Pombo Antunes, A. R. et al. Single-cell profiling of myeloid cells in glioblastoma across species and disease stage reveals macrophage competition and specialization. Nature Neuroscience 24, 595–610 (2021). 10.1038/s41593-020-00789-y

39 Friebel, E. et al. Single-Cell Mapping of Human Brain Cancer Reveals Tumor-Specific Instruction of Tissue-Invading Leukocytes. Cell 181, 1626-1642.e1620 (2020). 10.1016/j.cell.2020.04.055

40 Najem, H. et al. Central nervous system immune interactome is a function of cancer lineage, tumor microenvironment, and STAT3 expression. JCI Insight 7 (2022). 10.1172/jci.insight.157612

41 Chen, D. et al. CTLA-4 blockade induces a microglia-Th1 cell partnership that stimulates microglia phagocytosis and anti-tumor function in glioblastoma. Immunity 56, 2086-2104.e2088 (2023). 10.1016/j.immuni.2023.07.015

42 Sattiraju, A. et al. Hypoxic niches attract and sequester tumor-associated macrophages and cytotoxic T cells and reprogram them for immunosuppression. Immunity 56, 1825-1843.e1826 (2023). 10.1016/j.immuni.2023.06.017

43 Batchu, S., Hanafy, K. A., Redjal, N., Godil, S. S. & Thomas, A. J. Single-cell analysis reveals diversity of tumor-associated macrophages and their interactions with T lymphocytes in glioblastoma. Scientific Reports 13, 20874 (2023). 10.1038/s41598-023-48116-2

44 Bagley, S. J. et al. PD1 inhibition and GITR agonism in combination with fractionated stereotactic radiotherapy for the treatment of recurrent glioblastoma: A phase 2, multi-arm study. Journal of Clinical Oncology 41, 2004–2004 (2023). 10.1200/JCO.2023.41.16_suppl.2004

45 Ruiz-Moreno, C. et al. Harmonized single-cell landscape, intercellular crosstalk and tumor architecture of glioblastoma. bioRxiv, 2022.2008.2027.505439 (2022). 10.1101/2022.08.27.505439

46 Louveau, A. et al. CNS lymphatic drainage and neuroinflammation are regulated by meningeal lymphatic vasculature. Nat Neurosci 21, 1380–1391 (2018). 10.1038/s41593-018-0227-9

47 Badillo-Godinez, O. et al. Brain tumors induce immunoregulatory dendritic cells in draining lymph nodes that can be targeted by OX40 agonist treatment. Journal for ImmunoTherapy of Cancer 13, e011548 (2025). 10.1136/jitc-2025-011548

48 Ribas, A. et al. PD-1 Blockade Expands Intratumoral Memory T Cells. Cancer Immunology Research 4, 194–203 (2016). 10.1158/2326-6066.Cir-15-0210

49 Moseman, J. E., Rastogi, I., Jeon, D. & McNeel, D. G. PD-1 blockade employed at the time CD8+ T cells are activated enhances their antitumor efficacy. Journal for ImmunoTherapy of Cancer 13, e011145 (2025). 10.1136/jitc-2024-011145

50 Amoozgar, Z. et al. Targeting Treg cells with GITR activation alleviates resistance to immunotherapy in murine glioblastomas. Nat Commun 12, 2582 (2021). 10.1038/s41467-021-22885-8

51 Byrne, A. T. et al. Interrogating open issues in cancer precision medicine with patient-derived xenografts. Nat Rev Cancer 17, 254–268 (2017). 10.1038/nrc.2016.140

52 Golebiewska, A. et al. Side population in human glioblastoma is non-tumorigenic and characterizes brain endothelial cells. Brain 136, 1462–1475 (2013). 10.1093/brain/awt025

53 Ravi, V. M. et al. Spatially resolved multi-omics deciphers bidirectional tumor-host interdependence in glioblastoma. Cancer Cell 40, 639-655.e613 (2022). 10.1016/j.ccell.2022.05.009

54 Woroniecka, K. et al. T-Cell Exhaustion Signatures Vary with Tumor Type and Are Severe in Glioblastoma. Clin Cancer Res 24, 4175–4186 (2018). 10.1158/1078-0432.Ccr-17-1846

55 Chuprin, J. et al. Humanized mouse models for immuno-oncology research. Nat Rev Clin Oncol 20, 192–206 (2023). 10.1038/s41571-022-00721-2

56 Najem, H. et al. STING agonist 8803 reprograms the immune microenvironment and increases survival in preclinical models of glioblastoma. The Journal of Clinical Investigation 134 (2024). 10.1172/JCI175033

57 Liu, L. et al. Establishment and immune phenotyping of patient-derived glioblastoma models in humanized mice. Front Immunol 14, 1324618 (2023). 10.3389/fimmu.2023.1324618

58 Srivastava, R. et al. Development of a human glioblastoma model using humanized DRAG mice for immunotherapy. bioRxiv (2023). 10.1101/2023.02.15.528743

59 Quail, D. F. & Joyce, J. A. The Microenvironmental Landscape of Brain Tumors. Cancer Cell 31, 326–341 (2017). 10.1016/j.ccell.2017.02.009

60 Jangalwe, S., Shultz, L. D., Mathew, A. & Brehm, M. A. Improved B cell development in humanized NOD-scid IL2Rγnull mice transgenically expressing human stem cell factor, granulocyte-macrophage colony-stimulating factor and interleukin-3. Immunity, inflammation and disease 4, 427–440 (2016). 10.1002/iid3.124

61 Coughlan, A. M. et al. Myeloid engraftment in humanized mice: impact of granulocyte-colony stimulating factor treatment and transgenic mouse strain. Stem cells and development 25, 530–541 (2016). 10.1089/scd.2015.0289

62 Mathews, S. et al. Human Interleukin-34 facilitates microglia-like cell differentiation and persistent HIV-1 infection in humanized mice. Mol Neurodegener 14, 12 (2019). 10.1186/s13024-019-0311-y

63 Yaguchi, T. et al. Human PBMC-transferred murine MHC class I/II-deficient NOG mice enable long-term evaluation of human immune responses. Cellular & molecular immunology 15, 953–962 (2018). 10.1038/cmi.2017.106

64 Brehm, M. A. et al. Lack of acute xenogeneic graft-versus-host disease, but retention of T-cell function following engraftment of human peripheral blood mononuclear cells in NSG mice deficient in MHC class I and II expression. The FASEB Journal 33, 3137 (2019). 10.1096/fj.201800636r

65 Dobersalske, C. et al. Cranioencephalic functional lymphoid units in glioblastoma. Nature Medicine 30, 2947–2956 (2024). 10.1038/s41591-024-03152-x

66 Song, E. et al. VEGF-C-driven lymphatic drainage enables immunosurveillance of brain tumours. Nature 577, 689–694 (2020). 10.1038/s41586-019-1912-x

67 van Hooren, L. et al. Agonistic CD40 therapy induces tertiary lymphoid structures but impairs responses to checkpoint blockade in glioma. Nature Communications 12, 4127 (2021). 10.1038/s41467-021-24347-7

68 Cakmak, P. et al. Spatial immune profiling defines a subset of human gliomas with functional tertiary lymphoid structures. Immunity 10.1016/j.immuni.2025.09.018

69 Ramirez, C. F. A. & Akkari, L. Myeloid cell path to malignancy: insights into liver cancer. Trends in Cancer 11, 591–610 (2025). 10.1016/j.trecan.2025.02.006

70 Kung, H.-C., Zheng, K. W., Zimmerman, J. W. & Zheng, L. The tumour microenvironment in pancreatic cancer — new clinical challenges, but more opportunities. Nature Reviews Clinical Oncology (2025). 10.1038/s41571-025-01077-z

71 Bunse, L., Bunse, T., Kilian, M., Quintana, F. J. & Platten, M. The immunology of brain tumors. Science Immunology 10, eads0449 (2025). 10.1126/sciimmunol.ads0449

72 Brown, C. E. et al. Locoregional delivery of IL-13Rα2-targeting CAR-T cells in recurrent high-grade glioma: a phase 1 trial. Nature Medicine 30, 1001–1012 (2024). 10.1038/s41591-024-02875-1

73 Martins, T. A. et al. Enhancing anti-EGFRvIII CAR T cell therapy against glioblastoma with a paracrine SIRPγ-derived CD47 blocker. Nature Communications 15, 9718 (2024). 10.1038/s41467-024-54129-w

74 Garcia-Beltran, W. F. et al. Innate Immune Reconstitution in Humanized Bone Marrow-Liver-Thymus (HuBLT) Mice Governs Adaptive Cellular Immune Function and Responses to HIV-1 Infection. Frontiers in Immunology 12 (2021). 10.3389/fimmu.2021.667393

75 Hess, N. J., Brown, M. E. & Capitini, C. M. GVHD Pathogenesis, Prevention and Treatment: Lessons From Humanized Mouse Transplant Models. Frontiers in Immunology 12 (2021). 10.3389/fimmu.2021.723544

76 De Vleeschauwer, S. I. et al. OBSERVE: guidelines for the refinement of rodent cancer models. Nat Protoc 19, 2571–2596 (2024). 10.1038/s41596-024-00998-w

77 Montecino-Rodriguez, E. & Dorshkind, K. Use of Busulfan to Condition Mice for Bone Marrow Transplantation. STAR Protoc 1, 100159 (2020). 10.1016/j.xpro.2020.100159

78 Gerdes, M. J. et al. Highly multiplexed single-cell analysis of formalin-fixed, paraffin-embedded cancer tissue. Proc Natl Acad Sci U S A 110, 11982–11987 (2013). 10.1073/pnas.1300136110

79 Lindner, A. U. et al. An atlas of inter- and intra-tumor heterogeneity of apoptosis competency in colorectal cancer tissue at single-cell resolution. Cell Death Differ 29, 806–817 (2022). 10.1038/s41418-021-00895-9

80 Van der Walt, S. et al. scikit-image: image processing in Python. PeerJ 2, e453 (2014). 10.7717/peerj.453

81 Otsu, N. A Threshold Selection Method from Gray-Level Histograms. IEEE Transactions on Systems, Man, and Cybernetics 9, 62–66 (1979). 10.1109/TSMC.1979.4310076

82 Sauvola, J. & Pietikäinen, M. Adaptive document image binarization. Pattern Recognition 33, 225–236 (2000). 10.1016/S0031-3203(99)00055-2

83 Palla, G. et al. Squidpy: a scalable framework for spatial omics analysis. Nat Methods 19, 171–178 (2022). 10.1038/s41592-021-01358-2

