## Supplementary figures and tables for "Humanized glioblastoma patient-derived orthotopic xenografts recreate a locally immunosuppressed human immune ecosystem amenable to immunotherapeutic modulation"

bioRχiv

[1]NORLUX Neuro-Oncology Laboratory, Department of Cancer Research, Luxembourg Institute of Health, L-1210 Luxembourg, Luxembourg; [2]Faculty of Science, Technology and Medicine, University of Luxembourg, L-4367 Belvaux, Luxembourg; [3]Animal Facility, Department of Cancer Research, Luxembourg Institute of Health, L-4354 Esch-Sur-Alzette, Luxembourg; [4]Department of Physiology and Medical Physics, RCSI Centre for Systems Medicine, Royal College of Surgeons in Ireland University of Medicine and Health Sciences, D02 YN77 Dublin, Ireland; [5]Department of Infection and Immunity, Luxembourg Institute of Health, L-4354 Esch-Sur-Alzette, Luxembourg. [6]Neuro-Immunology Group, Department of Cancer Research, Luxembourg Institute of Health, L-1210 Luxembourg, Luxembourg; [7]Tumor Stroma Interactions, Department of Cancer Research, Luxembourg Institute of Health, L-1210 Luxembourg, Luxembourg

[#] Equal contribution

### SUPPLEMENTARY FIGURES

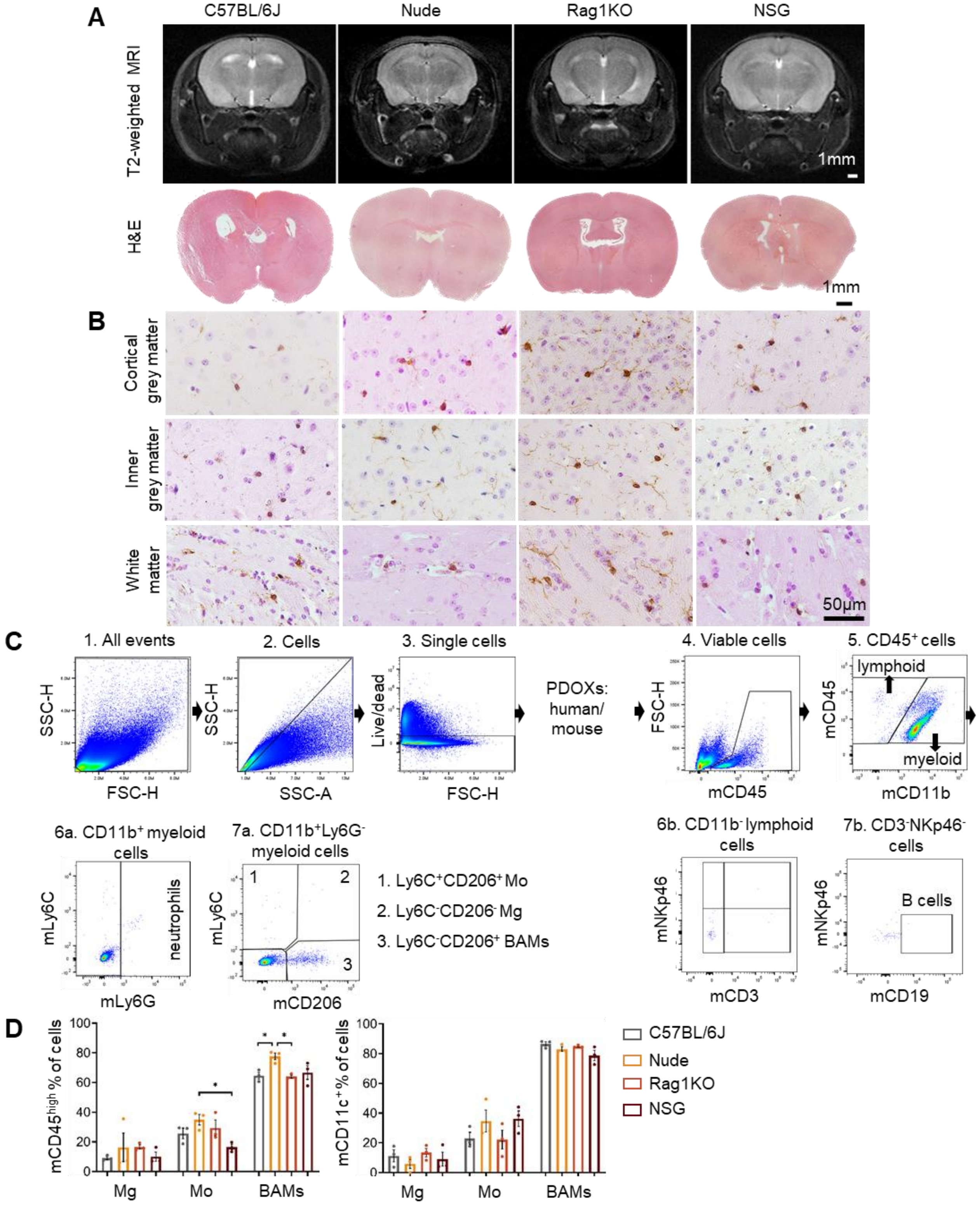


**Figure S1. Characterization of brain immune features in diverse mouse strains.** **A.** Representative T2-weighted MRI, and H&E staining of coronal sections of the entire brain from healthy 12-week-old mice in diverse mouse strains, scale bar: 1 mm. **B.** Representative Iba1 stainings in normal brains depicting myeloid cells. Images represent magnified areas in grey and white matter. Sections were co-stained with hematoxylin to visualize nuclei, scale bar: 50 µm. **C.** Flow cytometry gating strategy for profiling mouse-derived immune cells in the normal brain. Example is shown for a Nude CTR mouse: (1) Cells were distinguished from debris based on the Forward Scatter (FSC) and Side Scatter (SSC). (2) Cell aggregates were gated out based on their properties on the SSC area (SSC-A) versus height (SSC-H) dot plot. (3) Dead cells were excluded by their strong positivity for the dead cell marker. For PDOXs, discrimination between human and mouse cells is needed between step (3) and (4). (4) Mouse immune cells were recognized as mCD45$^{+}$ events (5) Lymphocytes were recognized as CD11b$^{-}$ population. Negative gating includes also CD11b$^{low}$ NK cells. CD11b$^{+}$ cells correspond to myeloid cells. Within myeloid cells (6a) CD11b$^{+}$Ly6G$^{+}$ cells represent neutrophils. (7a) Ly6C and CD206 allows distinguishing Ly6C$^{+}$CD206$^{-}$ Mo, Ly6C$^{-}$CD206$^{-}$ Mg and Ly6C$^{-}$CD206$^{+}$ BAMs. Within lymphoid cells: (6b) CD3 and NKp46 discriminates between T and NK cells respectively, (7b) CD3⁻NKp46⁻ cells are further interrogated for CD19⁺ B cell detection. **D.** Flow cytometry quantification highlighting baseline level expression of Ly6C⁻CD206⁻ Mg, Ly6C⁺CD206⁻ Mo and Ly6C⁻CD206⁺ BAMs for CD45 and CD11c retained in diverse mouse strains, one-way ANOVA with Tukey's HSD correction, comparing the mouse strains within each cell type (n = 3 mice/strain, mean ± SEM, *p<0.05).

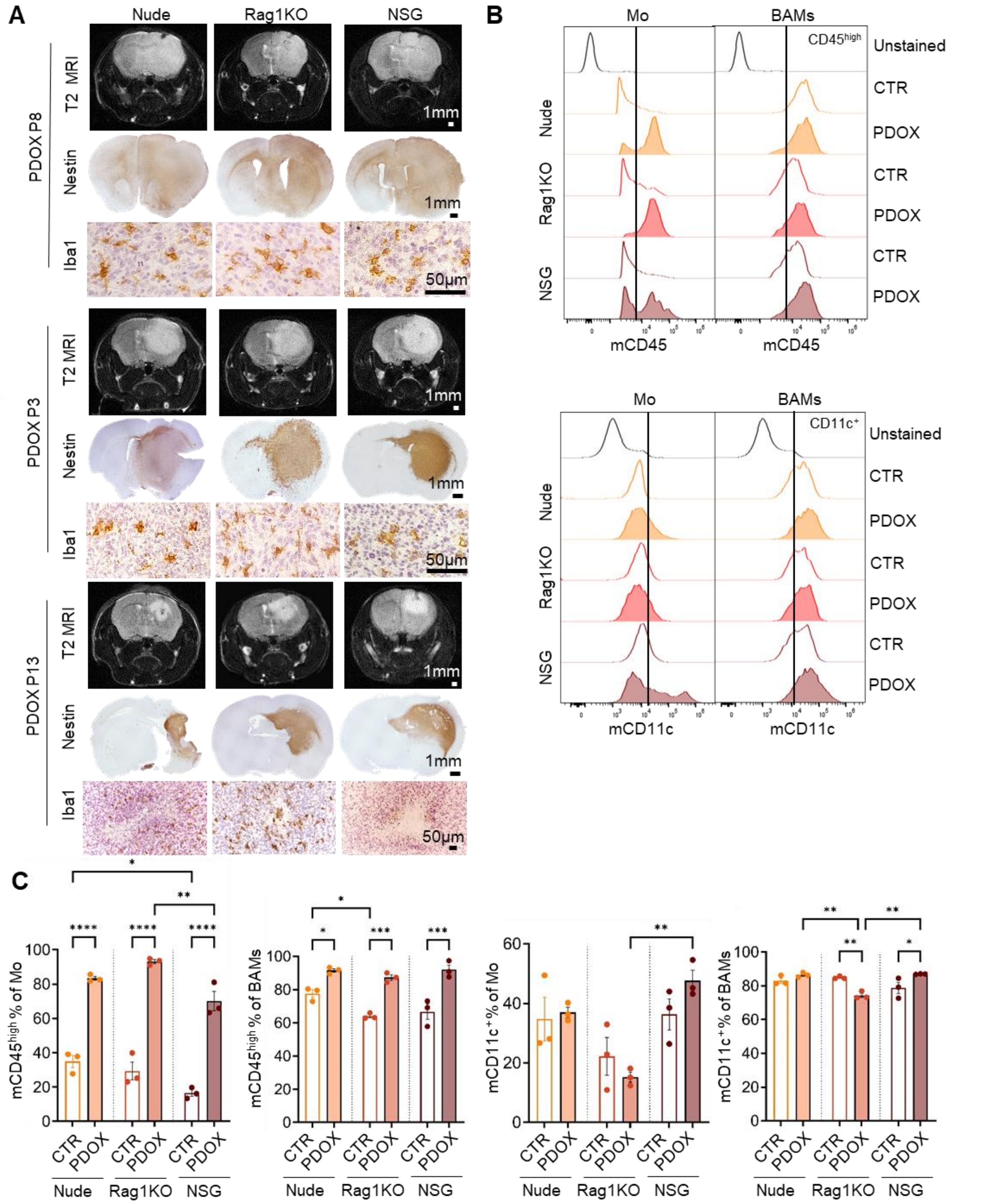


**Figure S2. Characterization of PDOX models derived across immunodeficient mouse strains. A.** Representative MRI and immunohistochemistry images in PDOX P8 (invasive), P3 (intermediate) and P13 (angiogenic) developed in Nude, Rag1KO and NSG mice. Nestin staining depicts human tumor cells in coronal sections of the entire brains, scale bar: 1 mm. Magnified Iba1 stainings show ameboid Mg within the cellular tumor areas, scale bar: 50 µm. **B-C.** Representative flow cytometry histograms and quantification showing increased CD45 and CD11c expression in Mo and BAMs within tumors of PDOX T188-implanted mouse brains compared to healthy control brains (CTR) across strains, one-way ANOVA with Tukey's HSD correction (n=3 mice/condition, mean ± SEM, *p<0.05, **p<0.01, ***p<0.001, ****p<0.0001).

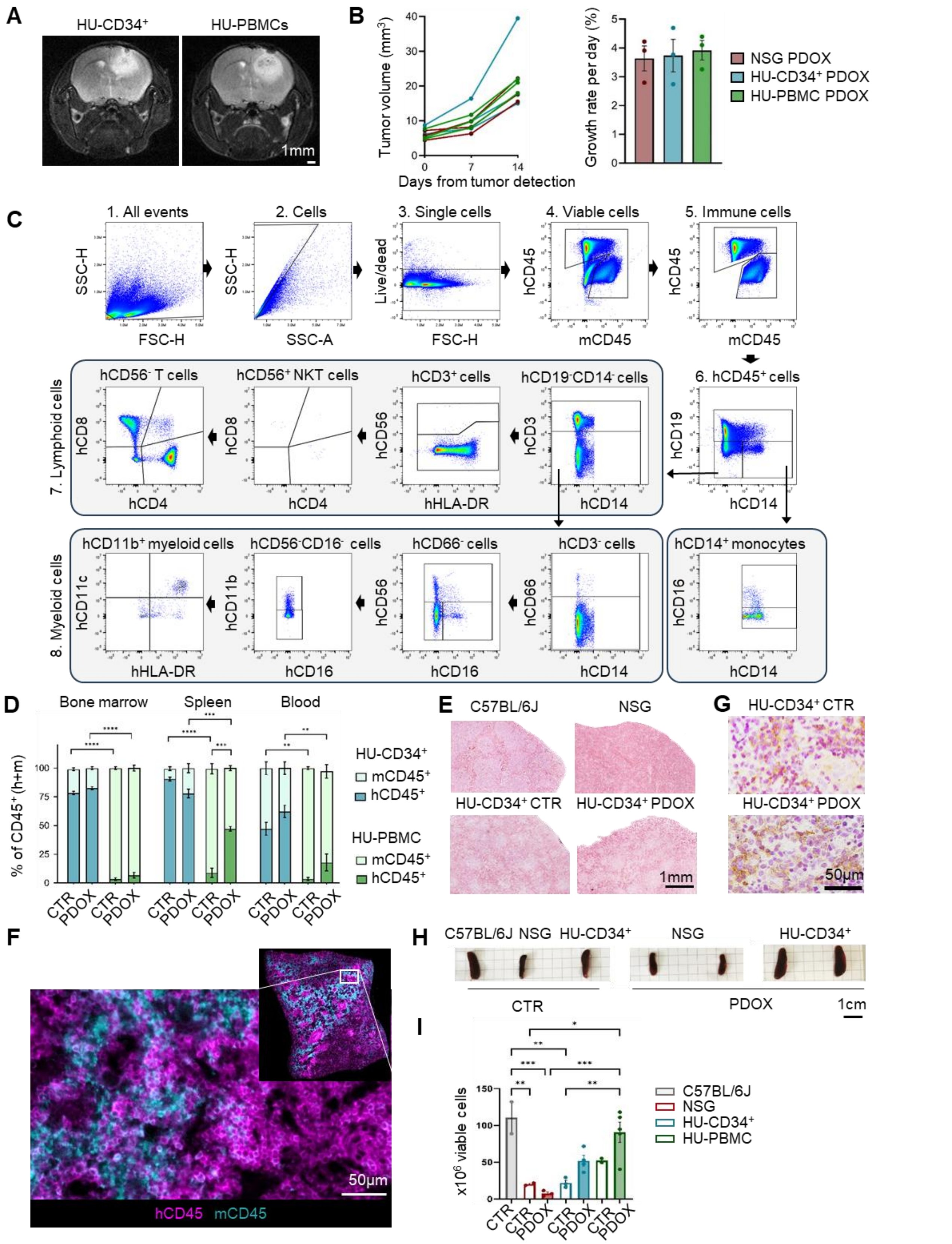


**Figure S3. Characterization of PDOX T188 and T158 tumor growth in HU-CD34+ and HU-PBMC mice. A.** MRI images of mouse brains in PDOX T158 developed in HU-CD34+ and HU-PBMC model, scale bar: 1 mm. **B.** MRI-based tumor volume (mm³) and growth rate per day quantification in NSG, HU-CD34+ and HU-PBMC PDOX T158 mice. Each connected dot line represents an individual mouse, one-way ANOVA with Tukey's HSD correction (n=3 mice/condition, mean ± SEM, ns = not significant). **C.** Flow cytometry gating strategy for profiling human immune cell types. Example is shown for spleen in HU-CD34+ T188 PDOX mouse: (1) Cells were distinguished from debris based on the Forward Scatter (FSC) and Side Scatter (SSC). (2) Cell aggregates were gated out based on their properties on the SSC area (SSC-A) versus height (SSC-H) dot plot. (3) Dead cells were excluded by their strong positivity for the dead cell marker (4) Mouse and human immune cells were recognized as $CD45^+$ events with species-specific antibodies. (5) Immune cells were quantified as a sum of $mCD45^+$ and $hCD45^+$ events. (6) Within human CD45 cells, classical monocytes were recognized as $CD14^+CD16^-$, B cells as $CD19^+$ events. (7) Lymphocytes were categorized as $CD3^+CD56^-$ T cells ($CD4^+$ and $CD8^+$) and $CD3^+CD56^+$ NKT cells (8) Myeloid cells were categorized into $CD3^-CD66^+$ neutrophils, $CD3^-CD66^-CD56^+$ NK cells and $CD3^-CD66^+CD56^-CD11b^+CD11c^+HLADR^+$ dendritic-like cells (DCs). **D**. Ratio between human ($hCD45^+$) and mouse ($mCD45^+$) immune cells in spleen, bone marrow, and blood in HU-$CD34^+$ (irradiated and busulfan combined) and HU-PBMC control mice (CTR) and PDOX T158. Statistical differences were assessed for hCD45 and mCD45 populations performing one-way ANOVA with Tukey's HSD correction (n=3 for CTR and PDOX mice/condition, mean ± SEM, **$p<0.01$, ***$p<0.001$, ****$p<0.0001$). **E**. H&E staining of magnified views from spleen sections of BL/6J, NSG, HU-$CD34^+$ CTR, and PDOX T188 mice, scale bar: 1 mm. **F.** Representative multiplex immunofluorescence of HU-$CD34^+$ PDOX T188 spleen, highlighting human CD45 (magenta) and mouse CD45 (cyan) immune cells, scale bar: 50 µm. **G**. hCD45 staining of spleen sections from HU-$CD34^+$ CTR and T188 PDOX mice, scale bar: 50 µm. **H.** Images of spleen in BL/6J, NSG and HU-$CD34^+$ CTR mice and NSG and HU-$CD34^+$ PDOXs depicting differences in size at endpoint, scale = 1 cm. **I.** Quantification of total viable cell numbers per spleen, one-way ANOVA with Tukey's HSD correction (n=2-3 for CTR, n=4-5 for T188 PDOX, mean ± SEM, *$p < 0.05$, **$p<0.01$, ***$p<0.001$).

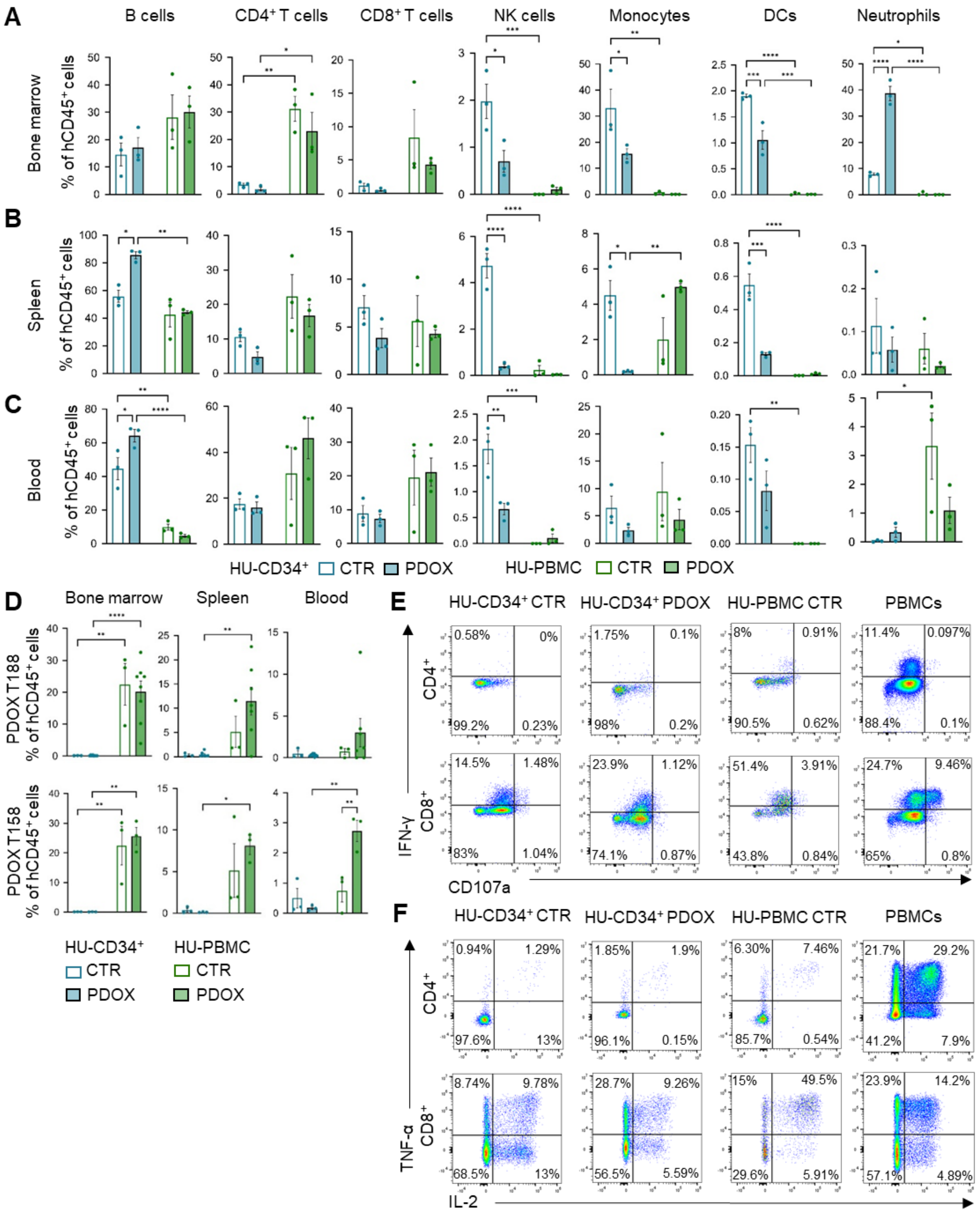


**Figure S4. Phenotypic profiling of human immune cells in bone marrow, spleen and blood in HU-CD34⁺ and HU-PBMC PDOXs.** **A-C.** Flow cytometry quantification of human lymphoid (B, T, NK cells) and myeloid populations (monocytes, DCs, neutrophils) in bone marrow (A), spleen (B), and blood (C) in HU-CD34⁺ and HU-PBMC CTR and PDOX T158 mice. Percentages indicate proportions of human CD45⁺ immune cells, one-way ANOVA with Tukey's HSD correction (n=3 mice/condition, mean ± SEM, *p<0.05, **p<0.01, ***p<0.001, ****p<0.0001). **D.** Quantification of CD4⁺CD8⁺ double-positive T cells in spleen, bone marrow, and blood across HU-CD34⁺ and HU-PBMC CTR and PDOX mice (T188 and T158 models). Percentages indicate proportions of total human CD45⁺ immune cells, one-way ANOVA with Tukey's HSD correction (n=3 for CTR and n=3-7 for PDOX mice/condition, mean ± SEM, *p<0.05, unpaired two-tailed Student's t-test). **E-F.** Flow cytometry dot plots of CD3⁺ T cells isolated from the spleens of HU-CD34⁺ and HU-PBMC CTR and PDOX T188 mice upon ex vivo stimulation with PMA/Ionomycin. Degranulation capacity (CD107a) and cytokine production (IFN-γ, A, IL-2, and TNF-α, B) were assessed. Stimulated human mature PBMCs are shown as reference.

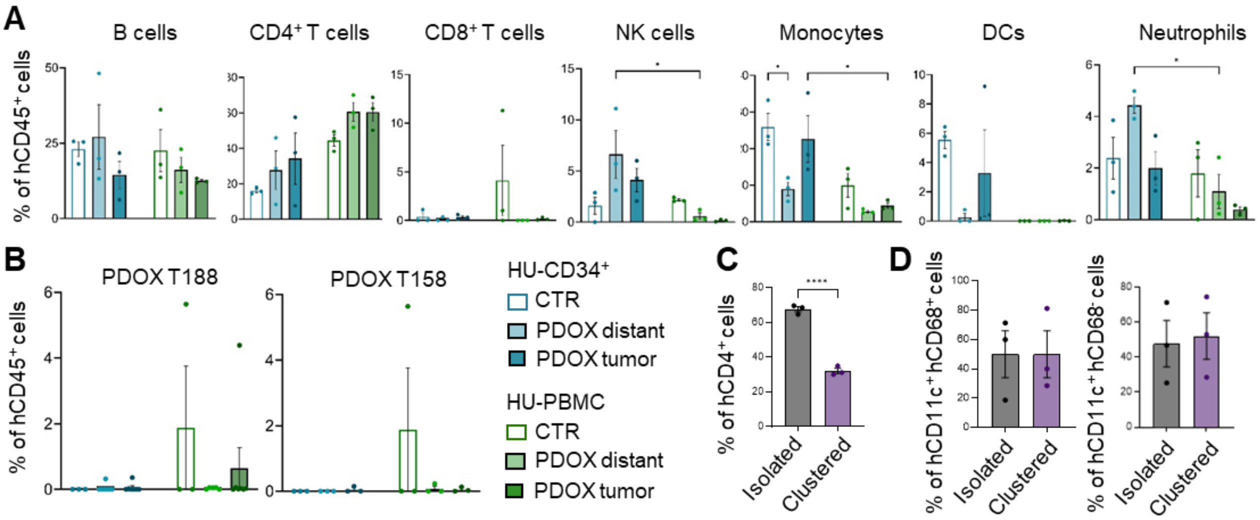


**Figure S5. Immune profiling and spatial distribution of infiltrating human immune cells in humanized GBM PDOXs. A.** Flow cytometry quantification of human lymphoid and myeloid populations in the brains of HU-CD34$^+$ and HU-PBMC mice. Percentages indicate proportions of human CD45$^+$ in control (CTR) and PDOX T158 mice ('tumor', and 'distant' areas), one-way ANOVA with Tukey's HSD correction (n=3 for CTR for PDOX mice/condition, mean ± SEM, *p<0.05). See gating strategy in **Fig S3C**. **B.** Quantification of CD4$^+$CD8$^+$ double-positive T cells in brains of HU-CD34$^+$ and HU-PBMC CTR and PDOX (T188 and T158 models). Percentages indicate proportions of human CD45$^+$ in control (CTR) and PDOX mice ('tumor', and 'distant' areas), one-way ANOVA with Tukey's HSD correction (n=3 for CTR and n=3-7 for PDOX mice/condition, mean ± SEM, not significant). **C-D.** Quantification of multiplex immunofluorescence of T188 HU-CD34$^+$ PDOX brains showing the proportion of isolated versus clustered CD4$^+$ T cells within the tumor (n=3, mean ± SEM, unpaired two-tailed Student's t-test,). Quantification of isolated versus clustered hCD4$^+$ T cells and hCD11c$^+$hCD68$^{+/-}$ myeloid cells within the tumor (n=3, mean ± SEM, unpaired two-tailed Student's t-test, **** p < 0.0001).

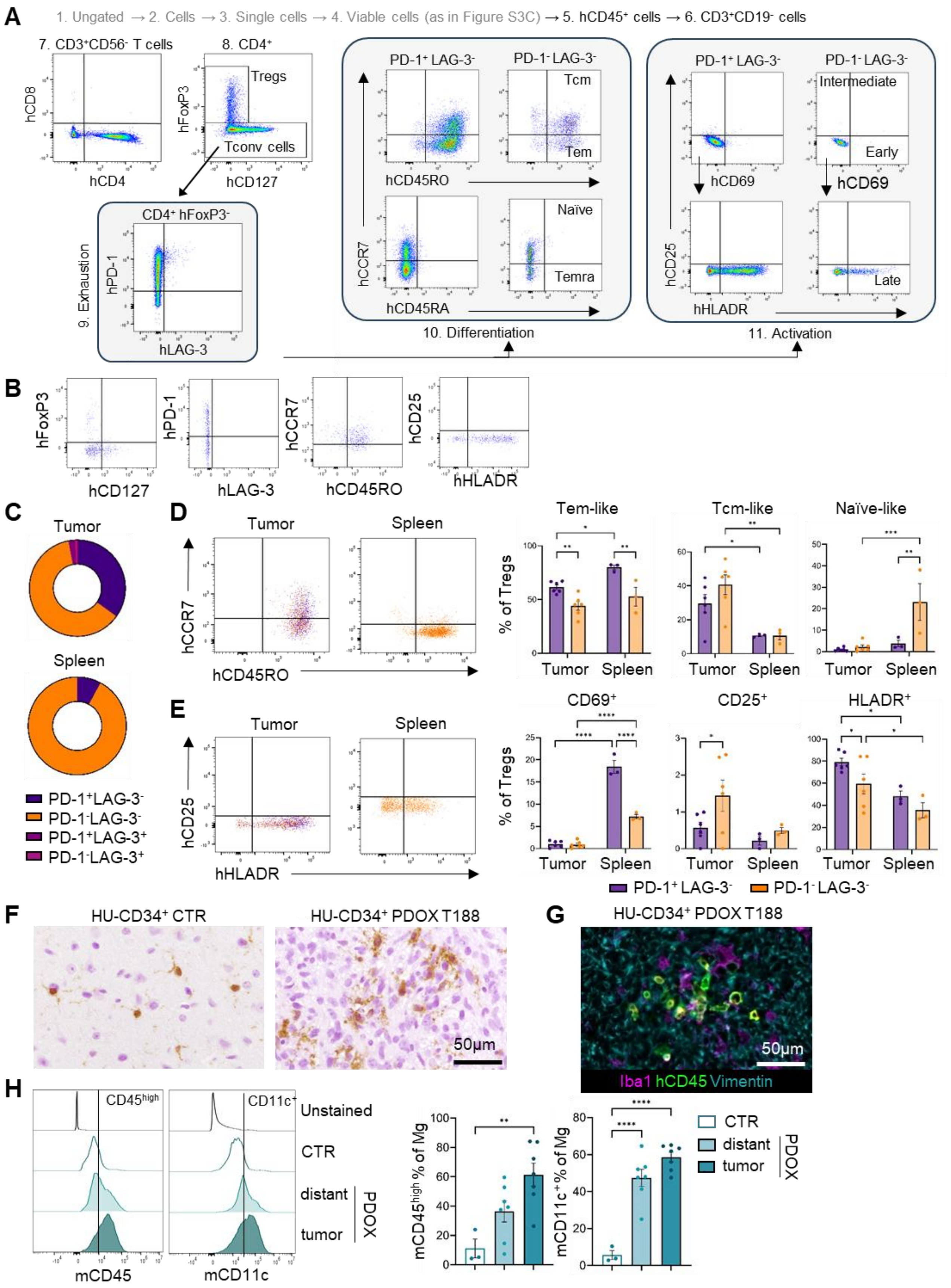


**Figure S6. Flow cytometry characterization of Treg phenotypes and murine TAMs in humanized GBM PDOXs. A.** Flow cytometry gating strategy for profiling human T cell phenotypic states. The gating strategy is consistent across organs, example is shown for tumor-bearing brain in PDOX T188 HU-CD34$^+$ mouse: (1-4 see gating strategy in **Fig S3C**) (5) Human lymphocytes were categorized as CD3$^+$ T or CD19$^+$ B cells (6) NKT cells were distinguished as CD56$^+$ events within CD3$^+$ cells. (7) CD56$^-$CD3$^+$cells were divided into CD8$^+$ and CD4$^+$ T cells for further functional annotation. (8) CD4$^+$ T cells were categorized into CD4$^+$FoxP3$^+$CD127$^{low}$ regulatory T cells (Tregs) or CD4$^+$FoxP3$^-$CD127$^{+/-}$ conventional T cells (Tconv). (9) Within each CD4$^+$ T cell subset, T cell exhaustion was identified via positivity for PD1 and/or LAG3. PD1$^+$LAG3$^-$

and PD1$^-$LAG3$^-$ subsets were analyzed separately for differentiation and activation to identify any potential differences associated to PD1 expression. (10) T cell differentiation was defined as effector memory-like (Tem, CD45RO$^+$CCR7$^-$), central memory-like (Tcm, CD45RO$^+$CCR7$^+$), effector memory CD45RA$^+$ T cells (Temra-like, CD45RO$^-$CD45RA$^+$CCR7$^-$), and naïve-like (CD45RO$^-$CCR7$^-$CD45RA$^+$) phenotypes. (11) T cell activation was investigated with early activation markers (CD69$^+$HLADR$^-$), intermediate (CD25$^+$) and late activation markers (CD69$^-$HLADR$^+$). **B.** Representative flow cytometry dot plots of CD4$^+$ T cell phenotypic states in HU-PBMC PDOX T188 tumor. Tregs were identified as CD4$^+$FoxP3$^+$CD127$^{low}$ cells. T cell exhaustion was assessed based on PD1 and LAG3 expression. T cell differentiation classified central memory-like (Tcm; CD45RO$^+$CCR7$^+$) and naïve-like (CD45RO$^-$CCR7$^+$) cells, and T cell activation was evaluated through HLADR and CD25 expression. **C.** PD1/LAG3-based exhaustion states in FoxP3$^+$ regulatory T cells (Tregs) cells in HU-CD34$^+$ PDOX T188 (mean, n = 6 for tumor, n = 3 for spleen). **D-E.** Representative flow cytometry dot plots display expression of (D) T cell differentiation and (E) activation markers in PD1$^+$LAG3$^-$ and PD1$^-$LAG-3$^-$ subsets in Tregs from tumor and spleen of HU-CD34$^+$ PDOX T188. Quantification of (D) T cell differentiation and (E) activation status in CD4$^+$ Tregs cells from HU-CD34$^+$ PDOX T188 brain tumors and spleens, PD1$^+$LAG3$^-$ exhausted and PD1$^-$LAG3$^-$ non-exhausted states are compared. T cell differentiation classified central memory-like (Tcm; CD45RO$^+$CCR7$^+$), effector memory-like (Tem; CD45RO$^+$CCR7$^-$) and naïve-like (CD45RO$^-$CCR7$^+$) cells. T cell activation was evaluated through CD69, CD25 and HLADR expression. For the gating strategy, see (A). Statistical comparisons were made between PD1$^+$LAG3$^-$ and PD-$^-$LAG3$^-$ subsets within tumors and spleen separately, as well as between tumor and spleen using two-way ANOVA (n=6 mice for PDOX tumor, n=3 for PDOX spleen, mean ± SEM, *p<0.05, **p<0.01, ***p<0.001, ****p<0.0001). **F.** Assessment of Mg morphology using Iba1 staining. Images represent magnified areas within cellular tumor regions in HU-CD34$^+$ CTR and PDOX T188 mice, scale bar: 50 µm. **G.** Representative multiplex immunofluorescence of HU-CD34$^+$ PDOX T188 brain, highlighting Iba1$^+$ myeloid cells (magenta), human CD45$^+$ immune cells (green), and Vimentin$^+$ tumor cells (cyan), scale bar: 50 µm. **H.** Representative flow cytometry histograms and quantification showing increased mouse CD45 and CD11c expression in Mg within tumors in PDOX T188 mouse brains ('tumor', and 'distant' areas) compared to healthy control brains (CTR) in HU-CD34$^+$ mice, confirming transition to TAMs, one-way ANOVA with Tukey's HSD correction (n=3 for CTR and n=7 for PDOX mice/condition, mean ± SEM, **p<0.01, ****p<0.0001).

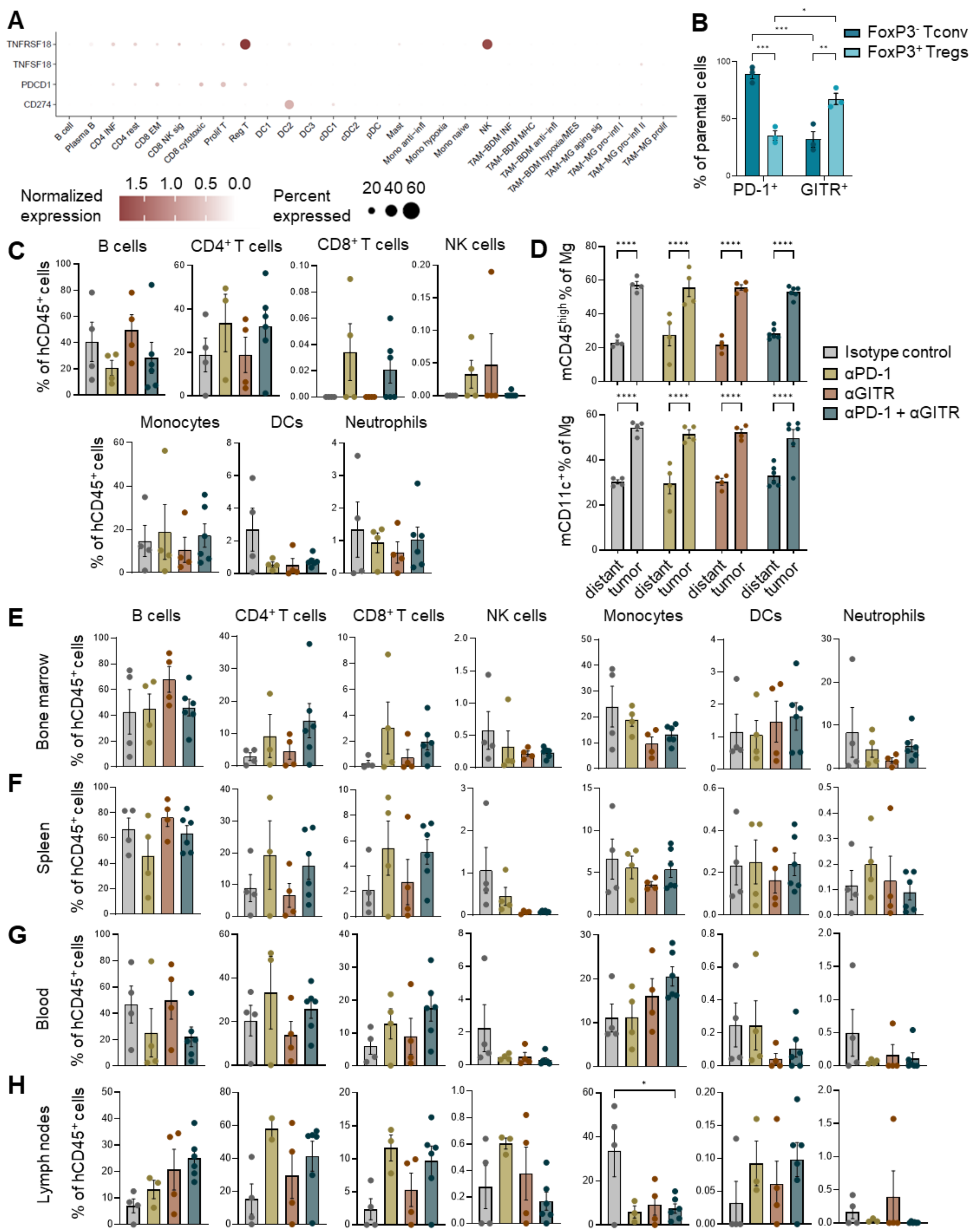


**Figure S7. GITR and PD-1 are expressed in GBM and humanized GBM PDOX models, and can be targeted systemically via antibody treatment.** **A.** Single-cell RNA-seq analysis showing gene expression of *TNFRSF18* (GITR), *TNFSF18* (GITR-L), *PDCD1* (PD-1) and *CD274* (PD-L1) in tumor-infiltrating immune cells from GBM patient samples. **B.** Flow cytometry quantification of PD-1 and GITR expression in $FoxP3^-$ T conventional (Tconv) and $FoxP3^+$ regulatory T cells (Tregs) from untreated HU-$CD34^+$ T188 PDOX mice, two-way ANOVA (n=3; mean ± SEM; *p<0.05, **p<0.01, ***p<0.001). **C–D.** Flow cytometry quantification of (**C**) human lymphoid and (**D**) myeloid subsets among human $CD45^+$ cells in the distant brain (left hemisphere) of HU-$CD34^+$ T188 PDOX mice across treatment arms, one-way ANOVA with Tukey's HSD correction (n=4 for isotype control, PD-1 and GITR monotherapy; n=6 for combination treatment; mean ± SEM; not significant). See gating strategy in **Fig. S3C**. **E.** Flow cytometry analysis of Mg within tumors of HU-$CD34^+$ T188 PDOX mice. Quantification of mouse CD45 and CD11c expression was performed in cells from 'tumor' and 'distant' regions across treatment arms, one-way ANOVA with Tukey's HSD correction (n=4 for isotype control, PD-1 and GITR monotherapy; n=6 for combination treatment; mean ± SEM; ****p<0.0001). **E-H.** Flow cytometry quantification of human lymphoid (B, T, NK cells) and myeloid populations (monocytes, DCs, neutrophils) in bone marrow (E), spleen (F), blood (G), and lymph nodes (H) in HU-$CD34^+$ T188 PDOX mice across treatment arms. Percentages indicate proportions of human $CD45^{+}$ immune cells, one-way ANOVA with Tukey's HSD correction (n=4 for isotype control, PD-1 and GITR monotherapy; n=6 for combination treatment; mean ± SEM; *p<0.05).

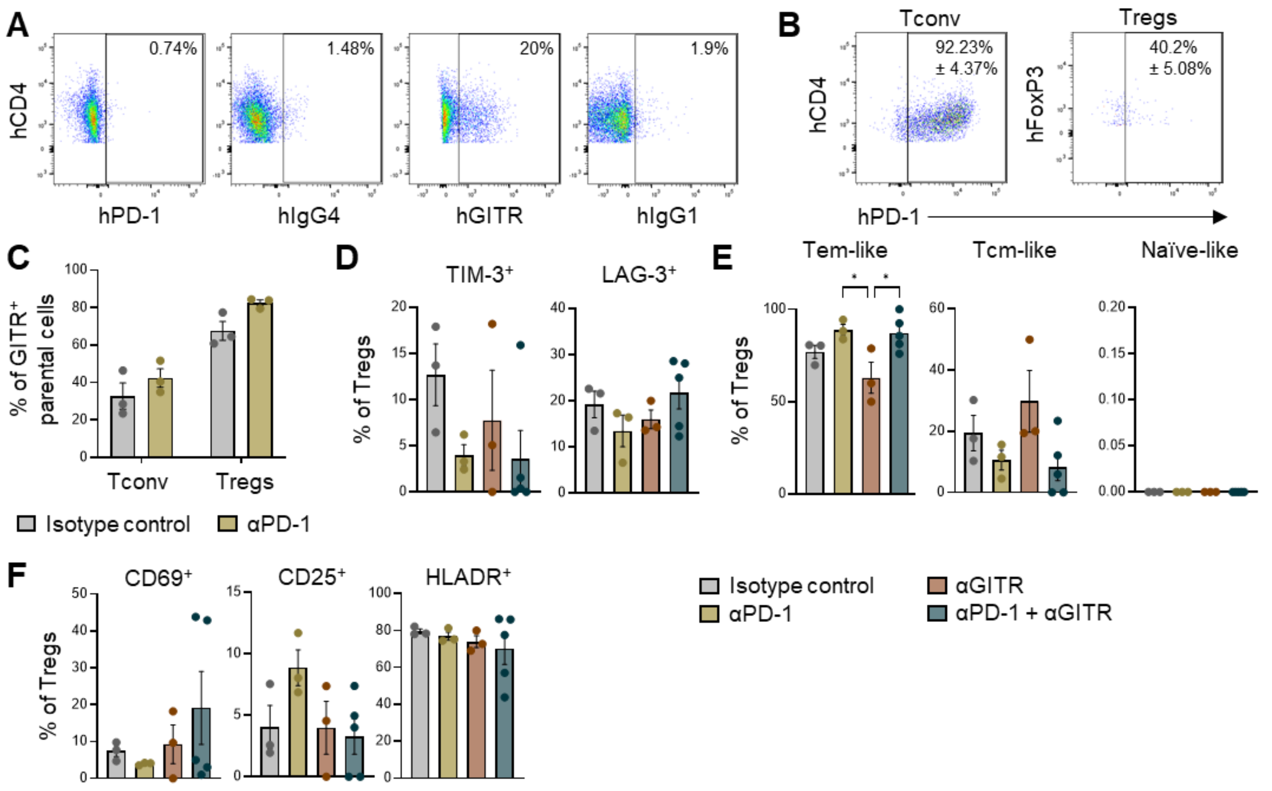


**Figure S8. Assessment of PD-1 and GITR detection and Treg phenotypes in HU-CD34+ PDOX mice. A.** Representative flow cytometry dot plots from a combination-treated HU-$CD34^{+}$ T188 PDOX mouse showing lack of detectable PD-1 signal using either an anti-PD-1 detection antibody or anti-IgG4 to reveal therapeutic antibody binding. Similarly, anti-GITR staining revealed partial signal, whereas anti-IgG1 did not detect therapeutic antibody-bound GITR. **B.** Representative dot plots from isotype control mice showing PD-1 expression in Tconv and Treg populations. Data represent mean ± SEM (n=3). **C.** Flow cytometry quantification of GITR expression in Tconv and Tregs from isotype control versus anti-PD-1-treated mice, unpaired, two-tailed Student's t-test (n=3 per group, not significant). **D-F.** Flow cytometry analysis of Tregs from HU-$CD34^{+}$ PDOX T188 across treatment arms. (**D**) Quantification of exhaustion markers TIM-3 and LAG-3. (**F**) Distribution of differentiation subsets, defined as effector memory-like (Tem; $CD45RO^{+}CCR7^{-}CD45RA^{-}$), central memory-like (Tcm; $CD45RO^{+}CCR7^{+}CD45RA^{-}$) and naïve-like ($CD45RO^{-}CCR7^{-}CD45RA^{+}$) cells. (**F**) Expression of activation markers CD69, CD25, and HLA-DR, one-way ANOVA with Tukey's HSD correction (n=3 for isotype control, PD-1 and GITR; n=5 for combination; mean ± SEM; *p<0.05).Gating strategy shown in **Fig. S5A**.

### SUPPLEMENTARY TABLES

**Table S1. Main characteristics of the PDOX models applied in the study.** The table depicts key patient characteristics (age, sex, tumor stage and diagnosis). In vivo characteristics and molecular profiles correspond to features detected in the preclinical models.

**Table S2. HLA genotyping of human immune cells applied in the study.**

**Table S3. List of antibodies used in the study.** Conjugated antibodies were applied for 3 multicolor panels: (i) mouse panel, Fig S1C; (ii) human panel, Fig S4C; (iii) T cell human panel, Fig S6E; (iv) cytokine panel, Fig S7D-E. CellDive multiplex immunofluorescence was carried out with custom conjugated antibodies or secondary Alexa dyes. *Flow cytometry test 106 cells/100µl.

**Table S4. CellDive imager configuration.**

**Table S1. Main characteristics of the PDOX models applied in the study.**

The table depicts key patient characteristics (age, sex, tumor stage and diagnosis). In vivo characteristics and molecular profiles correspond to features detected in the preclinical models.

| **Model** | **Patient** | | | **PDOX *in vivo* characteristics** | | | **PDOX molecular characteristics** | | | | | |
|---|---|---|---|---|---|---|---|---|---|---|---|---|
| PDOX model | Age at collection | Sex | Histological diagnosis | PDOX survival time (days) | Proliferation index in PDOX (%Ki67 +/- SD) | Detection by MRI | Copy number variations (CNV) | Glioma-specific gene mutations | Heidelberg DNA methylation classes | MGMT promoter methylation | GBM transcriptional subtypes | HLA profile |
| P3 | 64 | Male | GBM Grade IV | 42 +/- 5.2 | 54.7 +/- 3.7 | detectable, quantifiable | +[Chr7, 19p, 20q], -[1p36-1p34.1, 1q21.1-q44, -5p15.33-31, Chr9, Chr10, 11p15-14, 20p] --CDKN2A/B | PTEN (chr10:g.87957923del, ENSP00000361021.3:p.Asp236ThrfsTer20)<br>TP53 (chr17:g.7675085C>T, ENSP00000269305.4:p.Cys176Tyr) | GBM, IDHwt, RTK II | methylated | Classical | A*: 02:01:01, 02:01:01<br>B*: 27:05:02, 44:02:01<br>C*: 01:02:01, 05:01:01<br>DPA1*: 01:03:01, 01:03:01<br>DPB1*: 02:01:02, 04:01:01<br>DQA1*: 03:03:01, 03:03:01<br>DQB1*: 03:01:01, 03:01:01<br>DRB1*: 04:01:01, 04:08:01<br>DRB3*: No matches<br>DRB4*: 01:03:01<br>DRB5*: No matches |
| P8 | 64 | Female | GBM Grade IV | 52.5 +/- 3 | 38.3 +/- 10.8 | detectable, non quantifiable | ++EGFR, +[5p15.3-p12, 5q31.1-q35.3, Chr7, 8q24], -[ 6q21-q27, Chr10, 13q13.3-q34, 15q21.2-q23, 18q21.2-q22.4], --CDKN2A/B | EGFR (chr7:g.55154128G>A, ENSP00000275493.2:p.Ala289Thr)<br>MET (chr7:g.116771573G>T, ENSP00000317272.6:p.Ala954Ser)<br>PTEN (chr10:g.87961047_87961050del, ENSP00000361021.3:p.Thr319Ter);<br>ATRX (chrX:g.77663459C>G, ENSP00000362441.4:p.Ser1348Thr)<br>ATRX (chrX:g.77683638T>A, ENSP00000362441.4:p.Ser540Cys) | GBM, IDHwt, RTK I/II | methylated | Proneural | NA |
| P13 | NA | Female | GBM Grade IV | 35 +/- 2.5 | 42.9 +/- 10.8 | detectable, non quantifiable | +(Chr7, Chr19, Chr20), -(1p21.1-p31.2, 6q16.3-q21, Chr10, Chr13, 17q11-12), --CDKN2A/B | NF1 (chr17:g.31221849G>A, - )- likely not pathogenic | GBM, IDHwt, RTK II | methylated | Proneural | NA |
| T158 | 70 | Female | GBM Grade IV | 68 +/-1.6 | 35.2+/-3 | detectable, quantifiable | [++MDM2], +[Chr 7, Chr19, Chr20], -[Chr10, Chr15q], --CDKN2A/B | CDK6 (chr7:g.92615172T>A, ENSP00000265734.4:p.Ser317Cys)<br>NF1(chr17:g.31163302G>A, ENSP00000491431.1:p.Gly31Glu)-likely not pathogenic | GBM, IDHwt, RTK II | unmethylated | Classical | A*: No matches<br>B*: 07:02:01, 15:01:01<br>C*: 01:02:01, 07:02:01<br>DPA1*: 01:03:01, 02:01:04<br>DPB1*: 57:01, 117:01<br>DQA1*: No matches<br>DQB1*: 04:02:01<br>DRB1*: 15:01:01<br>DRB3*: 02:02:01<br>DRB4*: No matches<br>DRB5*: 01:01:01 |
| T188 | 68 | Male | GBM Grade IV | 63 +/- 3.9 | 37.1+/- 2.9 | detectable, quantifiable | [++EGFR], +[Chr7, Chr 19, Chr 20], -[1p36.23, 6p21.32, 6q27, Chr 10, 11q24.2], --CDKN2A/B | MDM4 (chr1:g.204549329A>C, ENSP00000356150.3:p.Lys374Gln)<br>PTCH1 (chr9:g.95459693C>T, ENSP00000332353.6:p.Val932Ile) | GBM, IDHwt, RTK II/I | unmethylated | Classical | A*: 02:01:01, 03:01:01<br>B*: No matches<br>C*: No matches<br>DPA1*: 01:03:01, 02:01:04<br>DPB1*: 04:01:01, 13:01:01<br>DQA1*: 01:03:01, 01:03:01<br>DQB1*: 06:03:01, 06:03:01<br>DRB1*: 13:01:01, 13:01:01<br>DRB3*: 02:02:01, 02:02:01<br>DRB4*: No matches<br>DRB5*: No matches |

**Table S2. HLA genotyping of human immune cells applied in the study.**

| Immune cell source type | Donor gender | Supplier | Catalog number | HU-MICE model applied | HLA profile | Relevant PDOX model |
|---|---|---|---|---|---|---|
| $CD34^+$ hematopoietic stem cells | NA | JAX | IHCM | $CD34^+$ HU-MICE (JAX, physical irradiation) | A*: 02:01, 03:01 | T188 |
| $CD34^+$ hematopoietic stem cells | Female | Lonza | 2C-101 | $CD34^+$ HU-MICE (in-house, busulfan chemical irradiation) | NA | T188, T158 |
| PBMCs | Female | Immunospot | CTL-CP1 | PBMC HU-MICE | A*: 02:01, 03:01<br>B*: 07:02, 15:01<br>C*: 03:04, 07:02<br>DPA1*: 01:03<br>DPB1*: 04:01<br>DQA1*: 01:02, 01:03<br>DQB1*: 03:02, 06:02<br>DRB1*: 04:01, 15:01<br>DRB4*: 01:01<br>DRB5*: 01:01 | T188, T158 |

**Table S3. List of antibodies used in the study.**

Conjugated antibodies were applied for 3 multicolor panels: (i) mouse panel, Fig S1C; (ii) human panel, Fig S4C; (iii) T cell human panel, Fig S6E; (iv) cytokine panel, Fig S7D-E.

CellDive multiplex immunofluorescence was carried out with custom conjugated antibodies or secondary Alexa dyes.

*Flow cytometry test 106 cells/100µl

| Antibody | Supplier | Catalog number | Reactivity | Concentration µl/test* | Multicolor panel |
|---|---|---|---|---|---|
| mCD11b PerCP5.5. | Biolegend | 101228 | mouse | 5 | i |
| mCD11c FITC | Biolegend | 117306 | mouse | 1.25 | i |
| mCD19 BV650 | Biolegend | 115541 | mouse | 5 | i |
| mCD206 APC | Biolegend | 141708 | mouse | 2.5 | i |
| mCD3 PE-Cy7 | Biolegend | 100320 | mouse | 2.5 | i |
| mCD45 BV605 | Biolegend | 103140 | mouse | 2.5 | i, ii, iii |
| mCD80 BV510 | Biolegend | 104741 | mouse | 2.5 | i |
| mF4/80 PE | Biolegend | 123110 | mouse | 5 | i |
| mLy6C Pacific Blue | Biolegend | 128014 | mouse | 0.5 | i |
| mLy6G BV785 | Biolegend | 127645 | mouse | 2.5 | i |
| mNKP46 PE-Cy5 | Biolegend | 137647 | mouse | 5 | i |
| hCD107a BV785 | Biolegend | 328643 | human | 5 | iv |
| hCD11b BV785 | Biolegend | 301346 | human | 5 | ii |
| hCD11c BV650 | Biolegend | 337238 | human | 5 | ii |
| hCD127 BUV737 | BD | 612795 | human | 5 | iii |
| hCD14 PE-Cy7 | Biolegend | 367112 | human | 5 | ii |
| hCD16 BV711 | Biolegend | 302044 | human | 0.6 | ii |
| hCD19 BB700 | BD | 566396 | human | 5 | ii, iii |
| hCD197 APC-Cy7 | Biolegend | 353211 | human | 5 | iii |
| hCD25 BV711 | Biolegend | 302635 | human | 5 | iii |
| hCD3 BV510 | Biolegend | 300448 | human | 5 | ii, iii |
| hCD4 BV605 | Biolegend | 300555 | human | 5 | iv |
| hCD4 PE-Cy5 | Biolegend | 300510 | human | 1 | ii, iii |
| hCD45 FITC | Biolegend | 368508 | human | 5 | ii, iii |
| hCD45RA BV750 | Biolegend | 304165 | human | 5 | iii |
| hCD45RO BUV395 | BD | 564292 | human | 5 | iii |
| hCD56 PE-CF594 | BD | 564849 | human | 0.6 | ii, iii |
| hCD66 BV421 | Biolegend | 392916 | human | 5 | ii |
| hCD69 BV650 | Biolegend | 310933 | human | 5 | iii |
| hCD8 AF700 | Biolegend | 344724 | human | 5 | iii |
| hCD8 APC | Biolegend | 301014 | human | 1.7 | ii |
| hFoxP3 BV421 | Biolegend | 320124 | human | 5 | iii |
| hFoxP3 PE | Biolegend | 320107 | human | 2.5 | iii |
| hGITR PE-Cy7 | Biolegend | 371224 | human | 5 | iii |
| hHLA-DR BV785 | Biolegend | 307641 | human | 5 | iii |
| hHLA-DR PE | Biolegend | 327008 | human | 5 | ii |
| hIFNy BV421 | Biolegend | 506537 | human | 1.7 | iv |
| hIgG BV421 | BD | 562581 | human | 5 | iii |
| hIgG PE-Cy7 | BD | 561298 | human | 5 | iii |
| hIgG1 BUV805 | BD | 753711 | human | 5 | iii |
| hIgG4 PE | SouthernBiotech | 9200-09 | human | 5 | iii |
| hIL-2 PE | Biolegend | 500306 | human | 1.7 | iv |
| hLAG-3 AF647 | Biolegend | 369303 | human | 5 | iii |
| hPD-1 BV421 | Biolegend | 329920 | human | 5 | iii |
| hTIM-3 BUV805 | Invitrogen | 368-3109-41 | human | 5 | iii |
| hTIM-3 PE-Cy7 | Biolegend | 345014 | human | 5 | iii |
| hTNFa AF700 | Biolegend | 502927 | human | 5 | iv |
| Human TruStain FcX™ (Fc Receptor Blocking) | Biolegend | 422302 | human | 5 | ii, iii, iv |
| TruStain FcX™ (anti-mouse CD16/32) | Biolegend | 101320 | mouse | 1 | i |
| LIVE/DEAD™ Fixable Blue Dead Cell Stain | ThermoFisher | L23105 | human/mouse | 1 | iii |
| LIVE/DEAD™ Fixable Near-IR | Invitrogen | L34975 | human/mouse | 1 | i, ii, iv |
| GFAP | Agilent | Z0334 | human/mouse | IHC:1/1000 | |
| hCD45 | Cell Signaling | 13917 | human | IHC:1/400 | |
| Iba1 | Biocare Medical | CP 290A | human/mouse | IHC:1/1000 | |
| Nestin | Abcam | AB6320 | human | IHC:1/500 | |
| CD11c (D3V1E) - AF647 | Cell Signaling | 42756BC | human | Cell Dive: 10µg/mL | |
| CD4 - AF555 | Abcam | ab280849 | human | Cell Dive: 50µg/mL | |
| CD45 (D9M8I) - AF750 | Cell Signaling | 16529BC | human | Cell Dive: 10µg/mL | |
| CD45RO (UCHL1) - AF488 | Biolegend | 304212 | human | Cell Dive: 5µg/mL | |
| CD68 (D4B9C) - AF555 | Cell Signaling | 23308BC | human | Cell Dive: 5µg/mL | |
| Iba1 | Wako | 019-19741 | human/mouse | Cell Dive: 10µg/mL | |
| PD-1 (D4W2J) - AF750 | Cell Signaling | 40948BC | human | Cell Dive: 0.5µg/mL | |
| PD-L1 (E1L3N®) - AF647 | Cell Signaling | 15005BC | human | Cell Dive: 20µg/mL | |
| SMA (α-smooth muscle actin) | Cell Signaling | 56856 | mouse | Cell Dive: 5µg/mL | |
| TIM-3 (D5D5R™) - AF647 | Cell Signaling | 78226S | human | Cell Dive: 15µg/mL | |

**Table S4. CellDive imager configuration.**

| Chanel band | Band (nm) | Center wavelegenth (nm) | (Bandwidth nm) |
|---|---|---|---|
| *Single-bandpass excitation filters* | | | |
| Blue | 379–401 | 390 | 22 |
| Green | 458–482 | 470 | 24 |
| Orange | 525.5–558.5 | 542 | 33 |
| Far red | 617–645 | 631 | 28 |
| Near IR | 710–750 | 730 | 40 |
| *Single-bandpass emission filters* | | | |
| Blue | 411–446 | 428.5 | 35 |
| Green | 495–512 | 503.5 | 17 |
| Orange | 574–599 | 586.5 | 25 |
| Far red | 664–690 | 677 | 26 |
| Near IR | 772–814 | 793 | 42 |
| *Polychroic beam splitter* | | | |
| Blue EX | 378–401 | 390 | 23 |
| Blue EM | 409–448 | 428.5 | 39 |
| Green EX | 457–483 | 470 | 26 |
| Orange EX | 526–558 | 542 | 32 |
| Orange EM | 571–603 | 587 | 16 |
| Far red EX | 617–645 | 631 | 28 |
| Far red EM | 660–695 | 677.5 | 35 |
| Near IR EX | 711–750 | 730 | 39 |
| Near IR EM | 767–820 | 793.5 | 53 |
